# Precise control of microtubule disassembly in living cells

**DOI:** 10.1101/2021.10.08.463668

**Authors:** Grace Y. Liu, Shiau-Chi Chen, Kritika Shaiv, Shi-Rong Hong, Wen-Ting Yang, Shih-Han Huang, Ya-Chu Chang, Hsuan Cheng, Yu-Chun Lin

## Abstract

Microtubules (MTs) are components of the evolutionarily conserved cytoskeleton, which tightly regulates various cellular activities. Our understanding of MTs is largely based on MT-targeting agents, which, however, are insufficient to dissect the dynamic mechanisms of specific MT populations due to their slow effects on the entire pool of MTs in cells. To address this limitation, we have used chemogenetics and optogenetics to disassemble specific MT subtypes by rapid recruitment of engineered MT-cleaving enzymes. Acute MT disassembly swiftly halted vesicular trafficking and lysosome dynamics. We also used this approach to disassemble MTs specifically modified by tyrosination and several MT-based structures including primary cilia, mitotic spindles, and intercellular bridges. These effects were rapidly reversed by inhibiting the activity or MT association of the cleaving enzymes. The disassembly of targeted MTs with spatial and temporal accuracy enables to uncover new insights of how MTs precisely regulate cellular architectures and functions.

## Introduction

Microtubules (MTs) are hollow tubes constructed from α- and β-tubulins and are present in all eukaryotic cells. They are involved in many cellular activities including intracellular trafficking, cell migration, cell division, cell polarity, signaling, and others^1,2^. To execute these functions, cells form several MT-based structures spatiotemporally such as primary cilia on the surface of G0 cells, intercellular bridges in the connected regions of two dividing cells during telophase, mitotic spindles in the cytosol of metaphase cells, and centrosomes^3–6^. Our understanding of MTs and these MT-based organelles is largely dependent on MT-targeting agents (MTAs) that perturb MT dynamics^7^. As MTs are vital for cellular physiology and mitosis, these MTAs have also been used in cancer chemotherapy^8^. In addition to their effects on MT dynamics, however, MTAs slowly disrupt MT structures and non-selectively perturb all MT subtypes. Several recent studies using photoswitchable MTAs and optogenetics to module MT dynamics or their ability to crosslink with actin filaments raise the possibility of precisely controlling MT activities^9–11^. However, none of these systems can directly disassemble specific MT subtypes and MT-based structures. We have developed a new and easy-to-use system that enables precise disassembly of targeted MT subtypes or specific MT-based structures under the control of chemical treatments or light illumination.

## Results

### Rapid translocation of proteins of interest onto MTs

The chemically inducible dimerization (CID) system has been used to manipulate cellular signaling and molecular composition over both space and time^12–14^. This is achieved by the rapid recruitment of proteins of interest (POIs) onto specific subcellular sites or their substrates^12^. Dimerization of FKBP (FK506-binding protein) and FRB (FKBP-rapamycin binding domain) that is triggered by a small chemical component, rapamycin, is one well-established CID system^12^. To rapidly recruit POIs onto MTs, we tagged FRB with a MT-binding sequence, EMTB (the MT-binding domain of ensconsin), and a cyan fluorescent protein (CFP) for visualizing its distribution (Fig. 1a). A linescan analysis showed that the resulting construct, EMTB-CFP-FRB, colocalizes with cytosolic MTs labeled by an antibody against α-tubulin (Supplementary Fig. 1). The addition of rapamycin to HeLa cells led to the rapid translocation of yellow fluorescent protein–tagged FKBP (YFP-FKBP) onto EMTB-CFP-FRB–labeled MTs as evidenced by an increased FRET (fluorescence resonance energy transfer) signal on MTs (T_1/2_ of translocation: 5.73 ± 0.54 sec; Fig. 1b,c; Supplementary Video 1). In summary, these results demonstrated that cytosolic POIs can be rapidly translocated onto MTs through inducible protein dimerization.

**Fig. 1.**
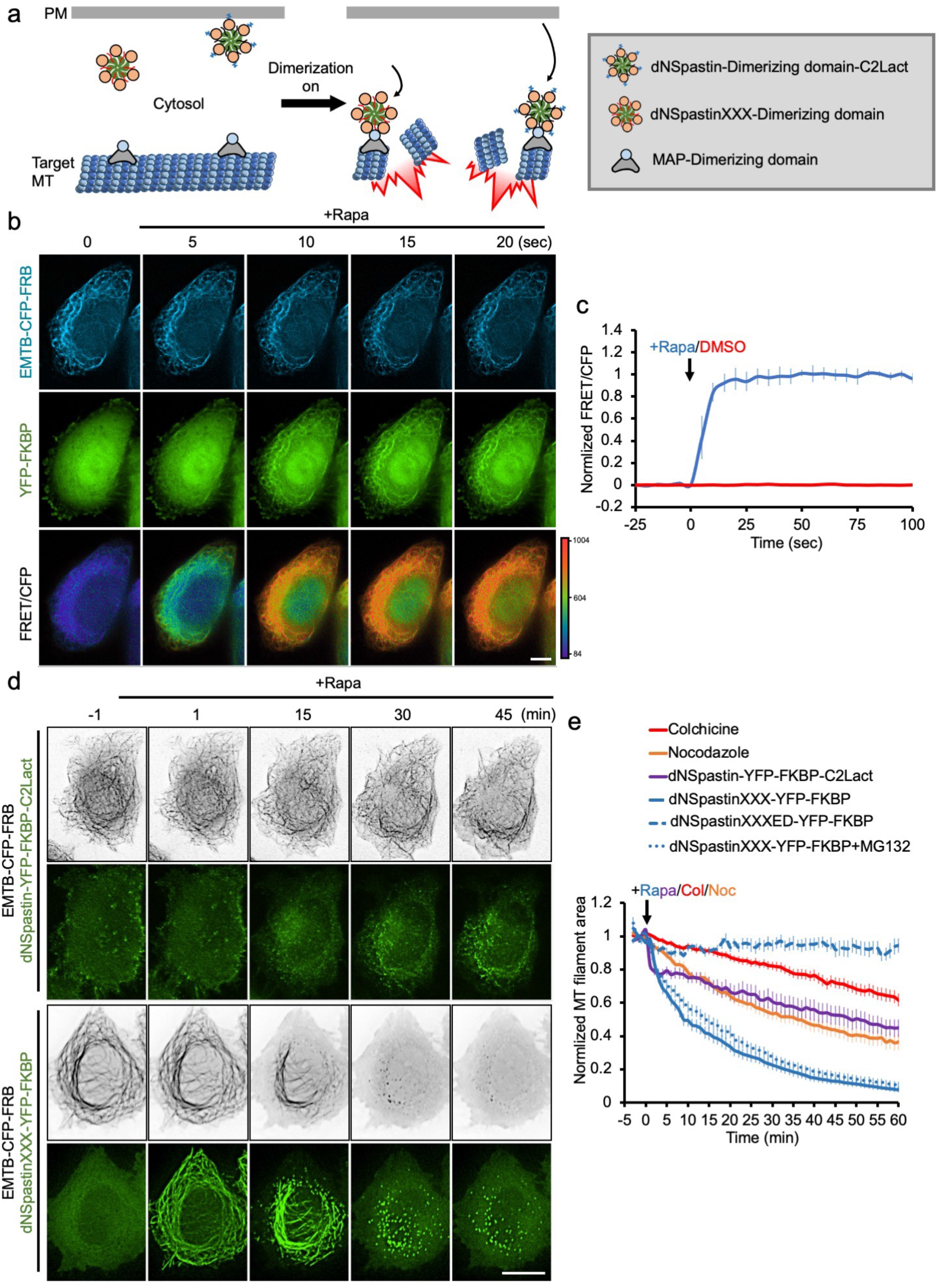
Recruitment of engineered Spastins onto MTs leads to their rapid disassembly. **a**. Schematic of the inducible MT disassembly system. One of the dimerization proteins is fused with one of two engineered Spastin enzymes, dNSpastinXXX and C2Lact-tagged dNSpastin, whereas the other is fused with MT-associated proteins (MAPs). Dimerization upon certain stimuli (e.g., chemical treatments or illumination) induces recruitment of engineered Spastins onto MAP-labeled MTs. Acute accumulation of Spastins on MTs rapidly induces disassembly of target MTs. **b**. HeLa cells co-transfected with EMTB-CFP-FRB and YFP-FKBP were treated with 100 nM rapamycin (Rapa). Addition of rapamycin rapidly translocated YFP-FKBP onto EMTB-CFP-FRB–labeled MTs and increased the FRET signal. Scale bar, 10 μm. **c**. The normalized intensity of FRET/CFP in cells before and after rapamycin (blue) and 0.1% DMSO (control; red) treatment. n = 6 and 10 cells in the rapamycin and DMSO group, respectively, from three independent experiments. **d**. The video frames of MTs after recruitment of the indicated enzymes onto MTs. HeLa cells co-transfected with the indicated constructs were treated with rapamycin (100 nM). Scale bar, 10 μm. **e**. The normalized MT filament area in cells with different MT disruption treatments. n = 30, 45, 19, 20, 14, and 6 cells in nocodazole (10 μM), colchicine (500 μM), dNSpastin-YFP-FKBP-C2Lact, dNSpastinXXX-YFP-FKBP, dNSpastinXXXED-YFP-FKBP, and dNSpastinXXX-YFP-FKBP with MG132 (50 μM) pre-treatment, respectively, from three independent experiments. Data are shown as the mean ± S.E.M.

### Engineering MT-severing enzymes for precise MT disruption

Spastin is a MT-severing enzyme that is ubiquitous in nearly all eukaryotic cells^15^. Consistent with previous study^16^, expression of full-length spastin (Spastin-YFP) and truncated spastin without the N-terminal 1–140 amino acids (dNSpastin-YFP) in HeLa cells removed significant amounts of cytosolic MTs, 47.45% and 42.86%, respectively, relative to YFP-transfected control cells (Supplementary Fig. 2). Thus the N-terminal fragment (1–140 amino acids) was dispensable for Spastin-mediated MT severing. To minimize the MT-cleaving reaction before recruitment of spastins onto MTs, we next attempted to disassociate dNSpastin from MTs by three strategies and also tagged spastin with YFP-FKBP: 1) removal of the MT-binding domain from dNSpastin^17^ (while retaining its catalytic domain; dNSpastinCD-YFP-FKBP); 2) mutation of key residues that are required for MT binding of Spastin (dNSpastinXXX-YFP-FKBP); and 3) tagging spastin with a plasma membrane–binding sequence, C2Lact^14^, to mislocate spastin to the cell cortex, where MTs are rarely found (dNSpastin-YFP-FKBP-C2Lact). We then characterized the enzyme activities of the above constructs before and after they were recruited onto MTs. dNSpastin-YFP-FKBP-C2Lact and dNSpastinXXX-YFP-FKBP showed low MT-severing activities before rapamycin treatment and robustly disrupted MTs after rapamycin treatment. dNSpastinCD-YFP-FKBP did not show severing activity (Supplementary Fig. 2). Live-cell imaging revealed that rapamycin treatment rapidly triggered the translocation of dNSpastin-YFP-FKBP-C2Lact (T_1/2_ = 55.44 ± 8.38 sec) and dNSpastinXXX-YFP-FKBP (T_1/2_ = 60.35 ± 5.30 sec) from the cytosol onto MTs (Supplementary Fig. 3; Supplementary Video 2). It was noteworthy that MT disruption triggered by dNSpastinXXX-YFP-FKBP upon rapamycin treatment (T_1/2_ = 11.05 ± 1.52 min) was much faster than that of dNSpastin-YFP-FKBP-C2Lact upon rapamycin treatment (T_1/2_ = 53.08 ± 8.08 min), and treatment of two common MTAs, nocodazole (10 μM; T_1/2_ = 54.02 ± 10.29 min), colchicine (500 μM; T_1/2_ = 94.28 ± 8.63 min; Fig. 1d,e; Extended data Fig. 1; Supplementary Videos 3, 4). Moreover, recruitment of dNSpastinXXX-YFP-FKBP onto MTs significantly removed more MT filaments as compared with others treatments (Extended data Fig. 1a,c). Trapping the enzyme-inactive dNSpastinXXXED-YFP-FKBP onto MTs showed no MT disruption, indicating that acute MT disruption is an enzyme-dependent event (Fig. 1e; Extended data Fig. 1; Supplementary Video 4). This inducible MT disruption occurs in other cell types (Supplementary Figs. 4, 5; Supplementary Videos 5, 6) and can also be controlled by rapamycin-orthogonal systems such as a gibberellin-based system with similar MT disruption kinetics (Supplementary Figs. 6, 7; Supplementary Video 7)^14^. Moreover, dNSpastinXXX-mediated MT disruption did not result in a change in the amount of MT mass when cells were pretreated with the proteasome inhibitor MG132, indicating that this event was not dependent on proteosome-mediated degradation (Fig. 1e; Extended data Fig. 2; Supplementary Video 8). Moreover, spastazoline, a spastin inhibitor^18^, swiftly reversed rapamycin-mediated acute MT disruption (Fig. 2; Supplementary Video 9). Intriguingly, nascent MTs from the fragments derived from dNspastinXXX-mediated cleavage assembled at a polymerization rate of 0.92 ± 0.04 μm/sec, which was slightly slower than the regrowth rate of 1.17 ± 0.07 μm/sec for MTs from centrosomes (Fig. 2c). Taken together, these results demonstrate that acute MT disruption occurs through MT filament disassembly instead of MT degradation. In summary, rapid recruitment of dNSpastinXXX-YFP-FKBP onto MTs efficiently triggered MT disassembly in living cells upon dimerization induction.

**Fig. 2.**
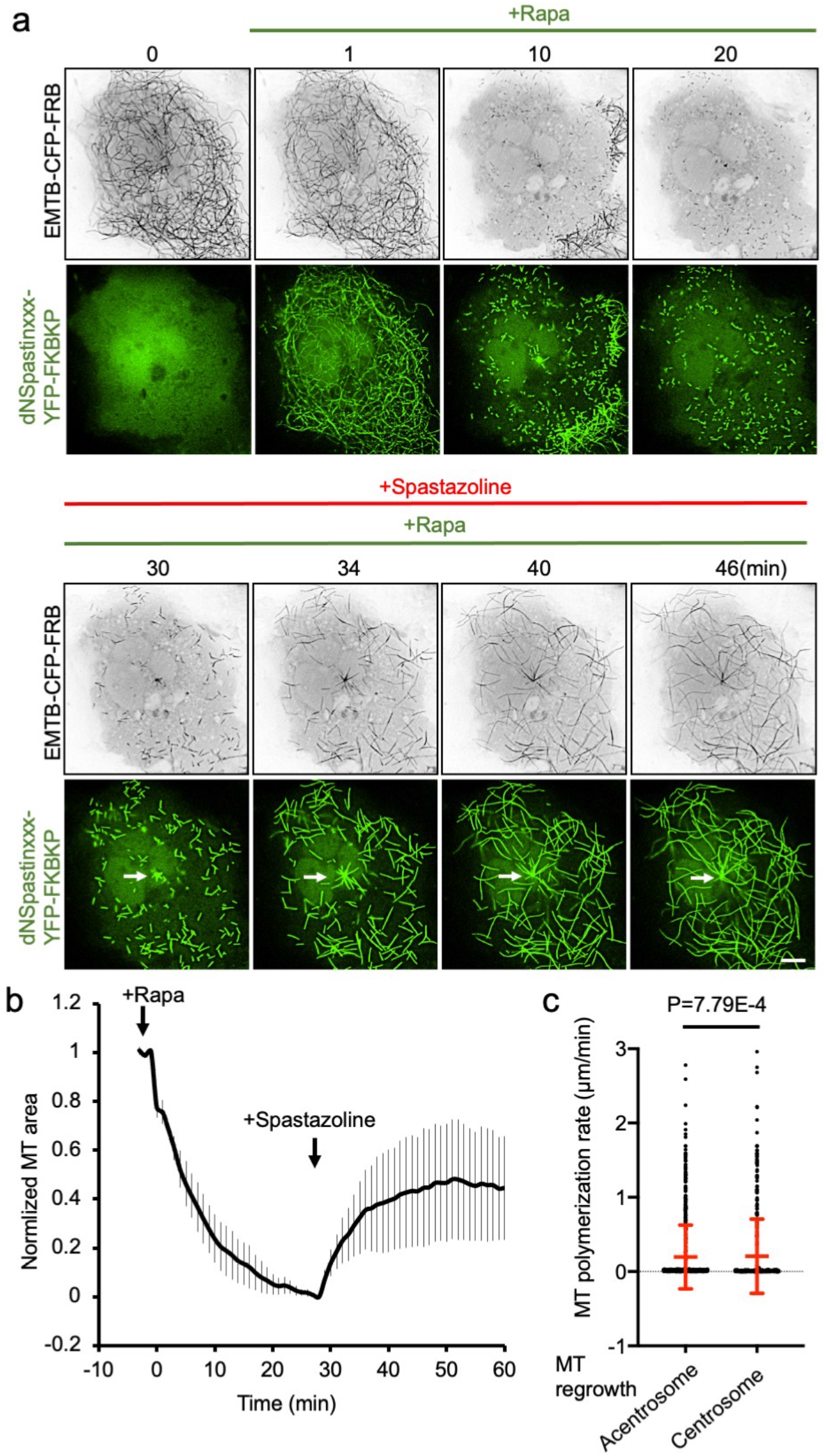
Inhibition of Spastin activity reverses MT disruption. **a**. HeLa cells co-transfected with EMTB-CFP-FRB (black) and dNSpastinXXX-YFP-FKBP (green) were pre-treated with rapamycin for 28 min to induce acute MT disassembly. Spastazoline (10 μM), a Spastin inhibitor, was then added to the cultures to halt our MT disruption system. Arrows indicate the centrosome-derived MTs. Scale bar, 10 μm. **b**. Normalized MT filament area for cells co-transfected and treated as in **a**. n = 3 cells from two independent experiments. Data are shown as the mean ± S.E.M. **c**. The polymerization rates of MTs regrowing from spastin-digested MT fragments in cytosol (acentrosome) and centrosomes. n = 147 and 74 MTs in the cytosol and centrosome group, respectively. Data (red) are shown as the mean ± SD. A Student’s t-test was performed to generate the indicated p-value.

### Acute MT disassembly inhibits vesicular trafficking and lysosome dynamics

We next evaluated whether acute MT disassembly functionally perturbs vesicle and organelle dynamics. To address this, a YFP-labeled post-Golgi vesicle marker, TGN38 (Trans-Golgi network integral membrane protein 38), and lysosome marker, LAMP3 (Lysosome-associated membrane glycoprotein 3), were used to observe the real-time dynamics of vesicles and lysosomes upon acute MT disassembly, respectively (Fig. 3 and supplementary Fig. 8). Acute MT disassembly attenuated the movement of post-Golgi vesicles and lysosomes in terms of moving distance and speed (Fig. 3 and supplementary Fig. 8; Supplementary Videos 10−13). Thus our system not only disrupts MT structure but also rapidly blocks vesicular trafficking and lysosome dynamics.

**Fig. 3.**
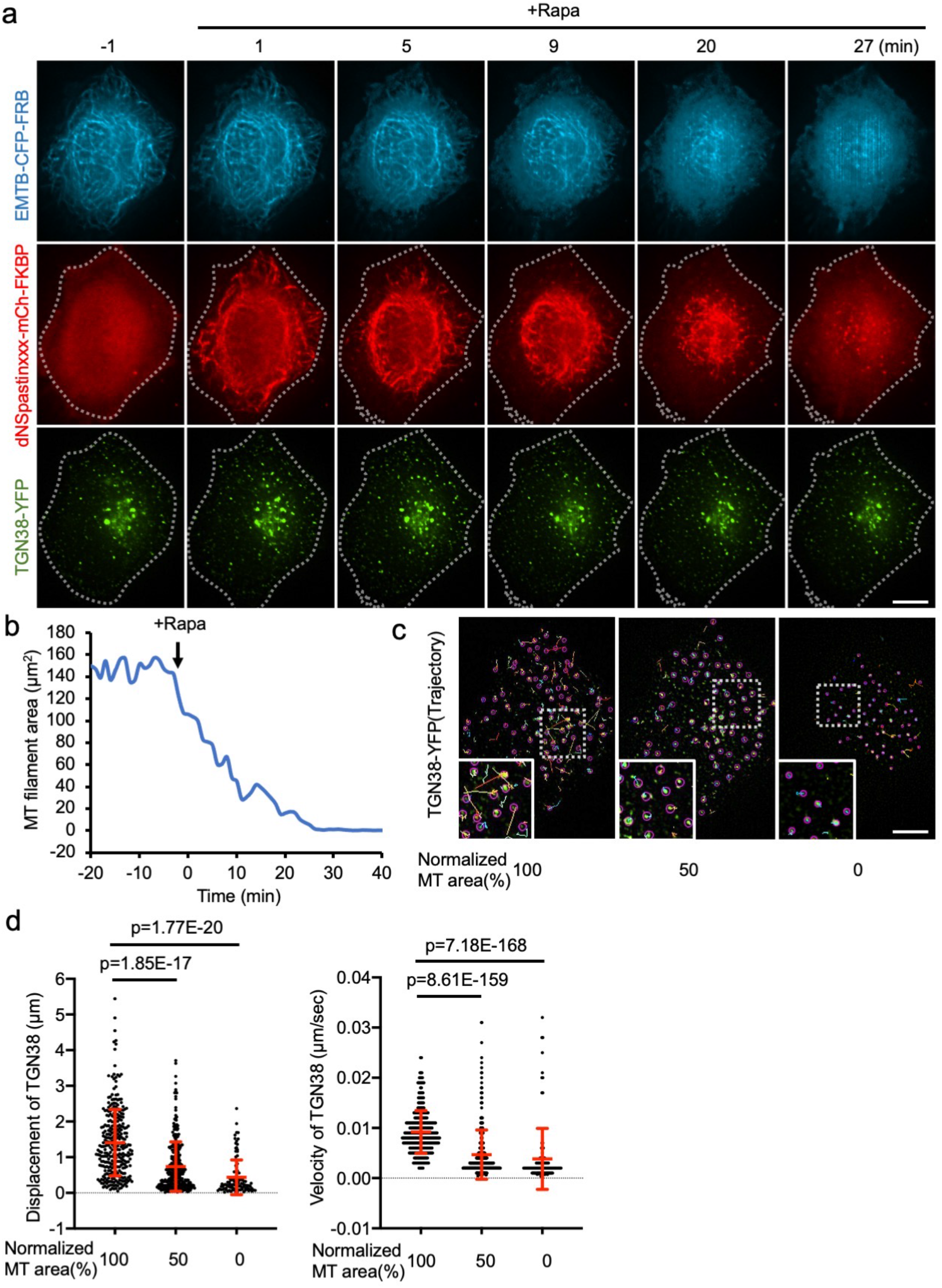
Acute MT disassembly attenuates vesicular trafficking. **a**. HeLa cells co-transfected with EMTB-CFP-FRB (blue), dNSpastinXXX-mCh-FKBP (red), and TGN38-YFP (**green**) were treated with rapamycin (100 nM) to induce MT disruption. Dotted lines indicate the cell boundary. Scale bar, 10 μm. **b**. The MT filament area in the cell shown in **a. c**. Trajectories of each TGN38-YFP−labeled vesicle at different levels of MT disruption. Insets show higher-magnification images of the regions indicated by the dotted-line boxes. Scale bar, 10 μm. **d**. The displacement (left) and velocity (right) of each labeled vesicle shown in **c**. n = 312, 319, and 111 vesicles in the MT 100, 50, and 0% group, respectively. Data (red) are shown as the mean ± SD. Student’s t-tests were performed to generate the indicated p-values.

### Rapid disruption of tyrosinated MTs

MTs undergo various post-translational modification (PTMs) for spatiotemporally regulating their properties and functions^1^. We next turned our efforts to precisely disrupt MTs with specific PTMs. Very recently, an elegant work has established a biosensor, A1AY1, that specifically target to tyrosinated MTs in living cells^19^. To apply it in our system, A1AY1 was tagged with FRB and a red fluorescent protein, TagRFP. Immunostaining results confirmed that the regulating construct (TagRFP-FRB-A1AY1) preferentially targeted to generic MTs and tyrosinated MTs but not to detyrosinated MTs (Fig. 4a, b). The addition of rapamycin rapidly recruited a cyan fluorescent protein, TagCFP, tagged FKBP (TagCFP-FKBP), onto TagRFP-FRB-A1AY1-bound MTs (T_1/2_ = 64.71 ± 16.08 sec; Supplementary Video 14). Moreover, recruitment of dNSpastinXXX-TagCFP-FKBP onto A1AY1-labeled MTs disrupted the A1AY1-positive MT filaments and significantly reduce ∼41.4% of tyrosinated MTs (Fig. 4e,f), indicating that, with specific tubulin PTMs biosensor against tyrosinated MTs, our system is able to precisely disassemble MTs modified with tyrosination.

**Fig. 4.**
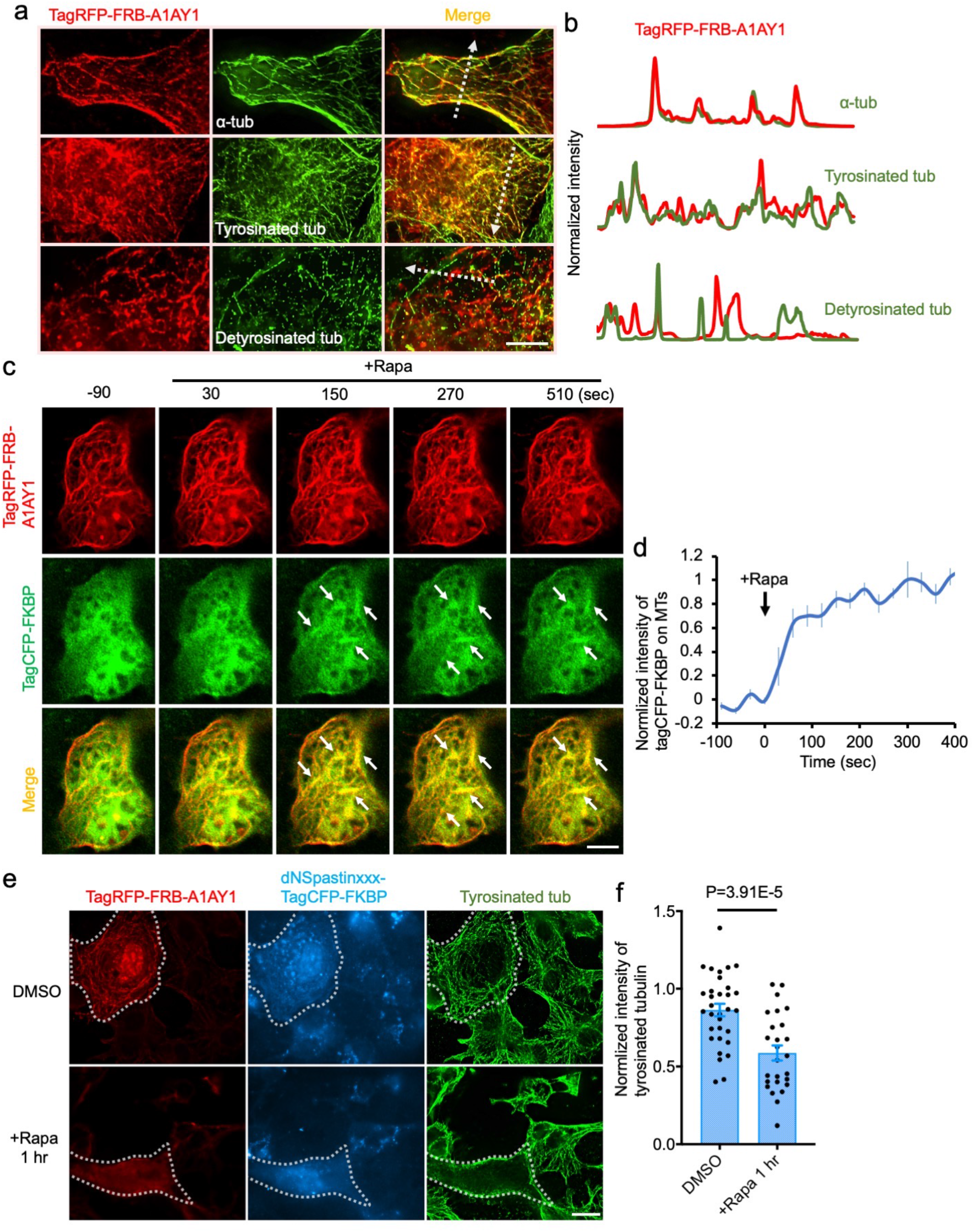
Disruption of tyrosinated MTs. **a**. COS7 cells transfected with TagRFP-FRB-A1AY1 (red) were labeled by anti-α-tubulin antibody (green; upper panel), anti-tyrosinated tubulin antibody (green; middle panel), or anti-detyrosinated tubulin antibody (green; lower panel), respectively. **b**. The intensity profiles of TagRFP-FRB-A1AY1 (red) and tubulin with the indicated PTMs (green) along dotted lines drawn in **a. c**. COS7 cells co-transfected with TagRFP-FRB-A1AY1 (red) and TagCFP-FKBP (green) were treated with rapamycin (100 nM). The addition of rapamycin rapidly recruits TagCFP-FKBP from cytosol onto A1AY1-labeled MTs (arrows). Scale bar, 10 μm. **d**. The normalized intensity of TagCFP-FKBP at A1AY1-labeled MTs upon rapamycin treatments. n=7 cells from 3 independent experiments. Data represent as mean ± S.E.M. **e**. COS7 cells co-transfected with TagRFP-FRB-A1AY1 (red) and dNSpastinXXX-TagCFP-FKBP (cyan) were treated with 0.1% DMSO or rapamycin (100 nM) for 1 hr and following by immunostaining with anti-tyrosinated tubulin antibody (green). Dotted lines highlight the transfected cells. Scale bar, 20 μm. **f**. The normalized intensity of tyrosinated MTs in TagRFP-FRB-A1AY1 and dNSpastinXXX-TagCFP-FKBP co-transfected cells upon 0.1% DMSO or rapamycin treatment for 1 hr. n=31 and 26 cells in DMSO and rapamycin treated groups, respectively, from three independent experiments. Data (blue) represent as mean ± S.E.M. Student’s t tests were performed with p values indicated.

### Rapid disruption of specific MT-based structures

In addition to cytosol MTs, many MT-based structures including primary cilia, centrosomes, mitotic spindles, and intercellular bridges regulate various cellular activities in a defined spatiotemporal manner^3–6^. We next tested whether recruitment of dNSpastinXXX onto these different MT-based structures can specifically disassemble them. A truncated MT-binding domain of microtubule associate protein 4 (MAP4m) preferentially bound to ciliary axonemes in G0 cells and shifts to mitotic spindles during metaphase and intercellular bridges during telophase (Supplementary Fig. 9)^13^. In G0 cells, addition of rapamycin rapidly trapped FKBP-tagged dNSpastinXXX on CFP-FRB-MAP4m−labeled axonemes in the cilia. Local accumulation of dNSpastinXXX on ciliary axonemes resulted in rapid disassembly of axonemes and primary cilia within ∼15 min (Fig. 5a, b; Extended data Fig. 3; Supplementary Video 15). Trapping enzyme-inactive dNSpastinXXXED-YFP-FKBP on ciliary axonemes did not perturb the ciliary structure (Extended data Fig. 3; Supplementary Video 15). Although axonemes are known as major skeletons of primary cilia, acute axoneme disruption was occasionally not associated with disassembly of the entire ciliary structure (2 out of 10 cells examined) and induced bulging and branched phenotypes in cilia (Supplementary Fig. 10; Supplementary Video 16), suggesting that other MT-independent factors may also contribute to the maintenance of ciliary structures. This may explain why rare cells paradoxically showed a ciliary membrane that did not have axonemes (Supplementary Fig. 11). In mitotic cells, translocation of dNSpastinXXX onto CFP-FRB-MAP4m−labeled mitotic spindles (Fig. 5c) and intercellular bridges (Fig. 5e) quickly disrupted these MT-based structures (Fig. 5c−f; Supplementary Videos 17, 18). Surprisingly, dNSpastinXXX did not disrupt centrosomes (or basal bodies in G0 cells) as shown by the images of cells that retained PACT (a conserved centrosomal targeting motif of pericentrin protein)^20^-labeled centrosomes after MT disruption treatment for 1 hr (Supplementary Fig. 12; Supplementary Video 19; the same phenotype is shown by dotted structures in Supplementary Videos 5, 6, 7, 9, 12, 15, and 16).

**Fig. 5.**
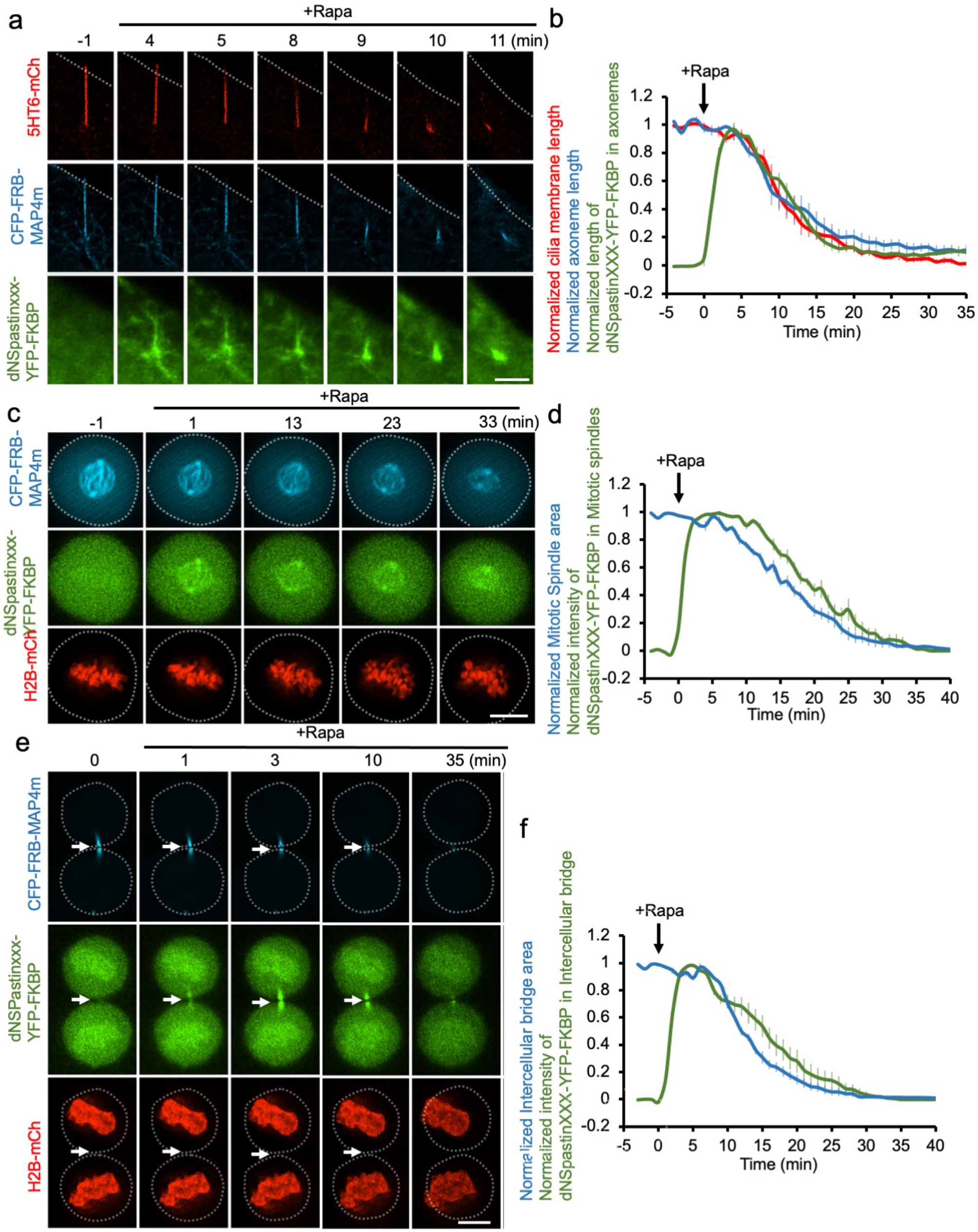
Rapid disruption of primary cilia, mitotic spindles, and intercellular bridges. **a**. NIH3T3 fibroblasts co-transfected with 5HT6-mCherry (5HT6-mCh; a ciliary membrane marker; red), CFP-FRB-MAP4m (blue), and dNSpastinXXX-YFP-FKBP (green) were serum starved for 24 hr to induce ciliogenesis. Ciliated cells were then treated with rapamycin (100 nM) to induce dNSpastinXXX-YFP-FKBP recruitment to axonemal MTs. Scale bar, 5 μm. **b**. The normalized length of axonemes (blue), primary cilia (red), and dNSpastinXXX-YFP-FKBP in cilia (green) upon rapamycin treatment. n = 6 cells from three independent experiments. **c**. HeLa cells co-transfected with H2B-mCherry (H2B-mCh; a chromosome marker; red), CFP-FRB-MAP4m (a marker of mitotic spindles; blue), and dNSpastinXXX-YFP-FKBP (green) were synchronized in metaphase and treated with rapamycin (100 nM) to induce dNSpastinXXX-YFP-FKBP recruitment to mitotic spindles. Scale bar, 10 μm. **d**. The normalized area of the mitotic spindle (blue) and intensity of dNSpastinXXX-YFP-FKBP in the mitotic spindle (green) upon rapamycin treatment. n = 6 cells from three independent experiments. **e**. HeLa cells co-transfected with H2B-mCherry (red), CFP-FRB-MAP4m (intercellular bridges; blue), and dNSpastinXXX-YFP-FKBP (green) were treated with rapamycin (100 nM) to induce dNSpastinXXX-YFP-FKBP recruitment to intercellular bridges. Arrows indicate the intercellular bridges. Scale bar, 10 μm. **f**. The normalized area of intercellular bridges (blue) and intensity of dNSpastinXXX-YFP-FKBP at intercellular bridges (green) upon rapamycin treatment. n = 5 cells from three independent experiments. Data are shown as the mean ± S.E.M. Dotted lines indicate the cell boundaries.

### Using light to spatiotemporally disrupt MTs

We next tried to use an optogenetic system to trigger MT disassembly in a sub-cellular region of interest during a specific period of time. Cryptochrome 2 (Cry2) and CIBN (N-terminal 170 amino acids of CIB1), two blue light−sensitive dimerizing partners^21^, were used in our MT-manipulating system. Cry2, which was tagged with the red fluorescent protein mCherry (mCh-Cry2), was rapidly translocated onto EMTB-YFP-CIBN−labeled MTs only in regions illuminated with light (T_1/2_ = 34.6 ± 7.58 sec) and dissociated from MTs to the cytosol when the light was off (T_1/2_ = 28.2 ± 4.88 sec; Fig. 6a, b; Supplementary Video 19). We then used light to control spastin-mediated MT disassembly in a spatially and temporally specific manner. dNSpastinXXX tagged with mCh-Cry2 was co-expressed with EMTB-YFP-CIBN in COS7 cells. MTs were labeled with SPY650-tubulin in these experiments. Local illumination with blue light robustly recruited dNSpastinXXX-mCh-Cry2 onto MTs, leading to the disassembly of MTs only in illuminated regions. The dNSpastinXXX-mCh-Cry2 reverted to cytosol and MT reassembly occurred when the cells were placed back in the dark (Fig. 6c, d; Supplementary Video 21). Recruitment of mCh-Cry2 without Spastin onto MTs by the same photostimulation procedure did not lead to MT disassembly, confirming that light-induced MT disassembly did not result from phototoxicity (Extended data Fig. 4; Supplementary Video 22).

**Fig. 6.**
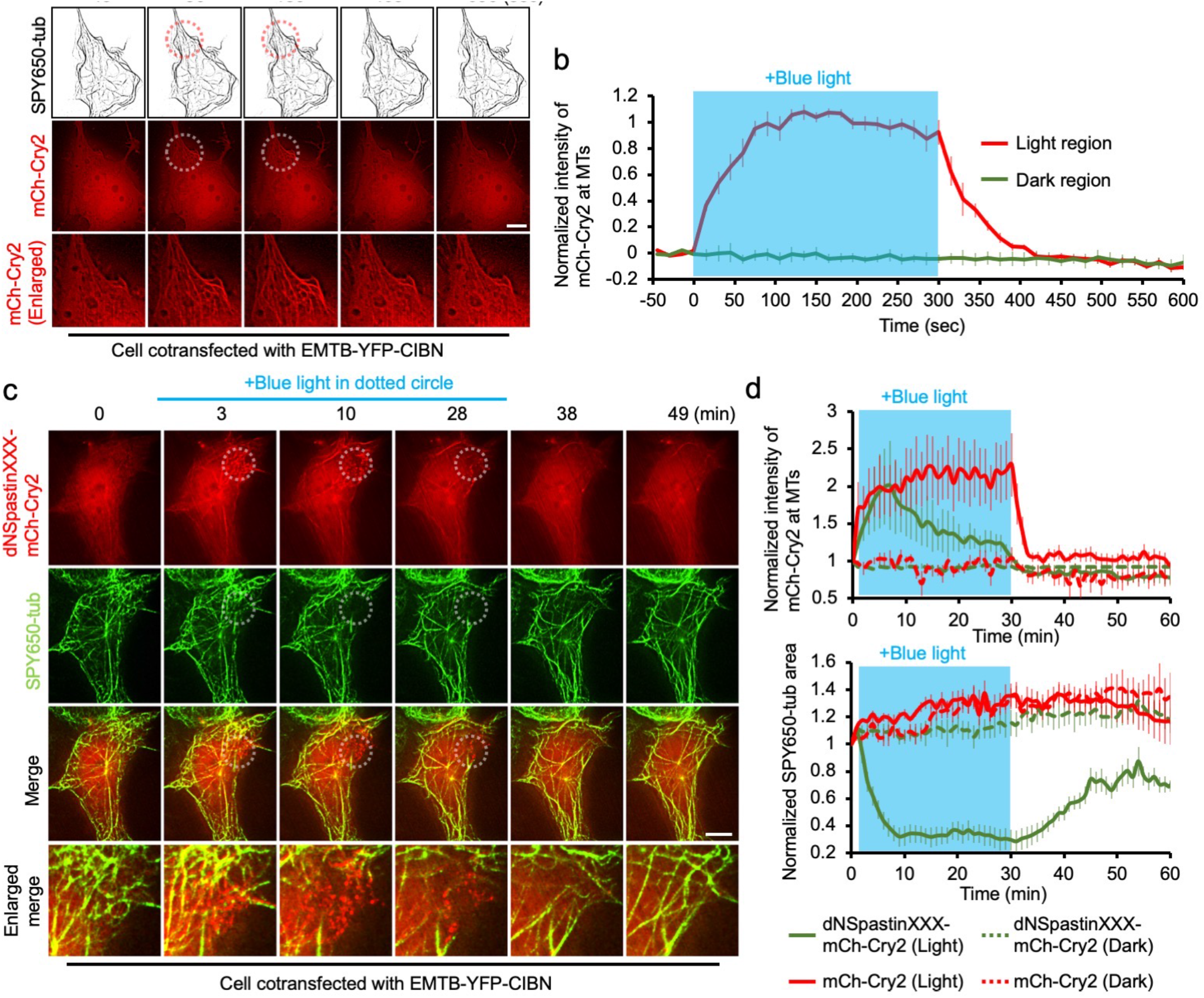
Using light to disassemble MTs in a reversible and location-specific manner. **a**. COS7 cells co-transfected with EMTB-YFP-CIBN and mCh-Cry2 (red) were incubated with SPY650-tubulin (SPY650-tub) to visualize MTs. The cells were illuminated by blue light within a specified region (indicated by the dotted circle) for the indicated time period. Scale bar, 10 μm. **b**. The normalized intensity of mCh-Cry2 at MTs in illuminated regions (red) and non-illuminated regions (green). n = 6 cells from three independent experiments. **c**. COS7 cells co-transfected with EMTB-YFP-CIBN and dNSpastinXXX-mCh-Cry2 were incubated with SPY650-tubulin to visualize MTs. The cells were illuminated by blue light as in **a**. Scale bar, 10 μm. **d**. The normalized intensity of dNSpastinXXX-mCh-Cry2 and mCh-Cry2 at MTs and the normalized area of SPY650-tubulin in illuminated regions and non-illuminated regions. n = 6 and 6 cells in dNSpastinXXX-mCh-Cry2 and mCh-Cry2 group, respectively, from three independent experiments. Data are shown as the mean ± S.E.M.

## Discussion

Overexpression of unmodified Spastin (SpastinFL) and dNSpastin depleted 47.45% and 42.86% of cytosolic MTs, respectively (Supplementary Fig. 2). Spastin-mediated cleavage of MTs was boosted by recruiting dNSpastin onto MTs with a 1-hr incubation of rapamycin which removed 74.75% of MTs (Supplementary Fig. 2). These results suggest a key role for MT association in regulating spastin-mediated MT cleaving. Previous *in vitro* and *in vivo* studies have demonstrated that tubulin glutamylation acts as a fine regulator to modulate MT binding and cleavage by Spastins, which supports this assumption^16,22^.

Spastin is involved in MT reorganization^23^. For example, Spastin preferentially localizes to axon branching points in neuronal cells to locally destabilize MTs and facilitate the elongation of branched MTs^24^. Our results showed that remnants of digested MTs derived from spastin-mediated severing were able to serve as platforms to extend nascent MTs when activity and MT association of spastin were inhibited (Figs. 2 and 6; Supplementary Videos 9 and 21). Several MT nucleation factors such as SSNA1 localize to axon branching sites and may cooperate with Spastin to initiate branched axons^25^. Whether MT nucleation factors accumulate on MT fragments upon this acute MT disassembly needs to be comprehensively evaluated to better understand the molecular mechanisms of MT remodeling.

The subcellular distribution of Spastin has been well studied, as it is preferentially enriched among cytosolic MTs, the mitotic spindle, and intercellular bridges^24,26^. Moreover, co-fractionation of Spastin with the centrosome marker γ-tubulin implies that Spastin may also be anchored to centrosomes/basal bodies^26^, and glutamylation on centrosomes may facilitate spastin-mediated severing^16^. To test whether Spastin is involved in MT severing in different subcellular regions, we recruited Spastin onto specific pools of MTs. Among all MT structures examined, the centrosome was the only one resistant to spastin-mediated MT severing (Supplementary Fig. 12; Supplementary Video 19). We suggest two explanations for this phenomenon. First, the unique triplet structure of centrosomal MTs may contribute to the centrosome’s ability to withstand spastin-mediated severing. A second possibility is that the anchoring orientation of EMTB-CFP-FRB may not allow the spastin-mediated severing reaction at centrosomes. Experiments to evaluate the effect of Spastin on centrosomes that lack MT triplets^27^ and that use different centrosomal-targeting proteins will be useful for addressing these possible mechanisms.

In addition to providing insights into how Spastin modulates MTs, our acute MT disassembly system is a powerful tool for distinguishing MT-dependent and -independent mechanisms and for dissecting causal relationships between MTs of interest and cellular events. This precise MT manipulation system also represents a fundamental step in developing new strategies to treat various diseases such as tumors and neurodegeneration and developmental disorders in which MT-mediated events have gone awry.

## Acknowledgements

We thank Dr. Carsten Janke (Institut Curie) for the Spastin constructs. This study was supported by the Ministry of Science and Technology (MOST), Taiwan (MOST grant numbers 109-2636-B-007-003 and 108-2638-B-010-001-MY2 to Y.C.L.).

## Contributions

G.Y.L., K.S., and Y.C.L. designed and conducted the experiments. G.Y.L., S.C.C., K.S., and Y.C.C. generated DNA constructs. S.R.H performed western blotting analysis. S.C.C., W.T.Y., and Y.C.C. conducted the photostimulation experiments. H.C. analyzed the dynamics of vesicles and lysosomes. S.H.H. conducted cell synchronization. Y.C.L., G.Y.L., and W.T.Y. wrote the manuscript.

## Ethics declarations

The patent for acute MT disassembly developed in this study is pending.

## Data availability

The data that support the findings of this study are available from the corresponding author upon reasonable request.

## Figure legends

**Supplementary Fig. 1.**
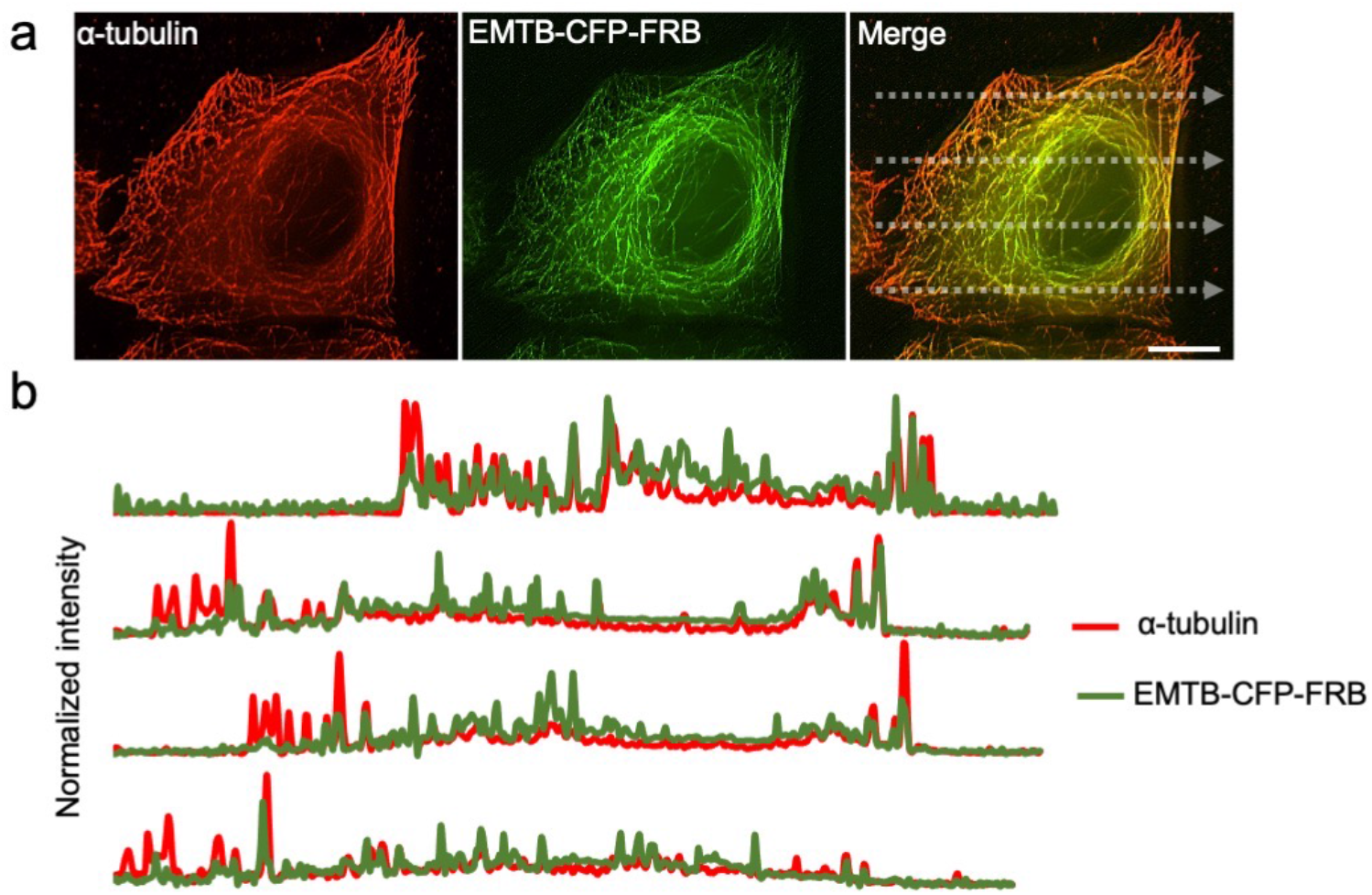
EMTB-CFP-FRB localizes to cytosolic MTs. **a**. HeLa cells expressing EMTB-CFP-FRB (green) were fixed and stained with anti-α-tubulin antibody (red). Scale bar, 10 μm. **b**. The normalized intensity profiles of α-tubulin (red) and EMTB-CFP-FRB (green) along the dotted lines drawn in **a**.

**Supplementary Fig. 2.**
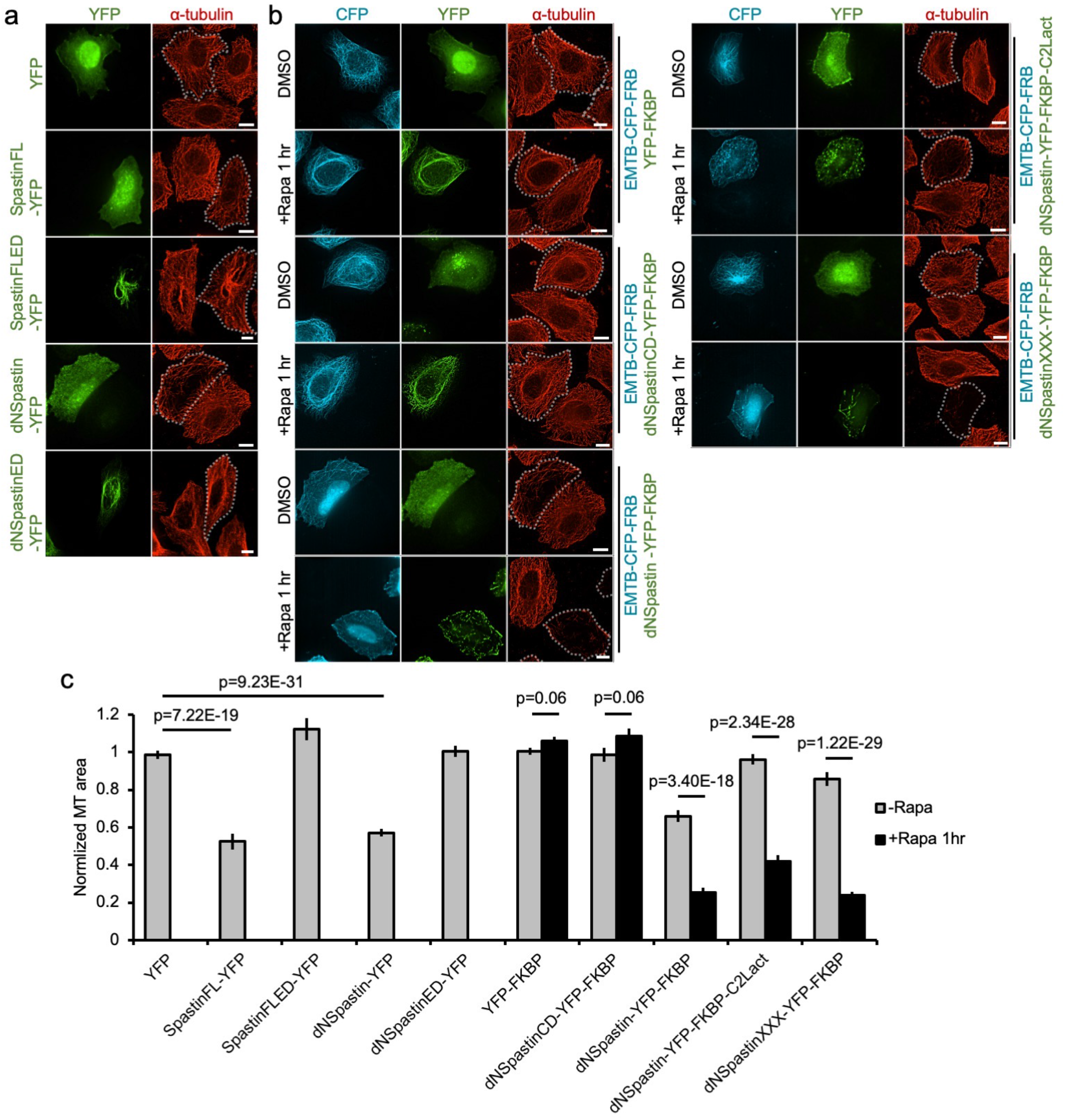
The MT-severing activity of engineered Spastin enzymes. **a**. HeLa cells transfected with YFP-tagged full-length spastin (SpastinFL-YFP; green) and truncated spastin (dNSpastin-YFP; green) were fixed and stained with the α-tubulin antibody (red). YFP alone and an ED (enzyme dead) form of each enzyme (SpastinFLED and dNSpastinED) are negative controls. Scale bar, 10 μm. **b**. HeLa cells co-transfected with EMTB-CFP-FRB and the indicated constructs were treated with either 0.1% DMSO or 100 nM rapamycin (Rapa) for 1 hr and then were fixed and stained with anti-α-tubulin antibody. Scale bar, 10 μm. Dotted lines highlight transfected cells. **c**. The normalized intensity of α-tubulin in cells expressing the indicated construct with 0.1% DMSO or rapamycin treatment. n = 103, 39, 33, 123, 41, 175, 144, 88, 79, 82, 90, 134, 104, 81, and 97 cells from left to right; three to five independent experiments. Data are shown as the mean ± S.E.M. Student’s t-tests were performed, with p-values indicated.

**Supplementary Fig. 3.**
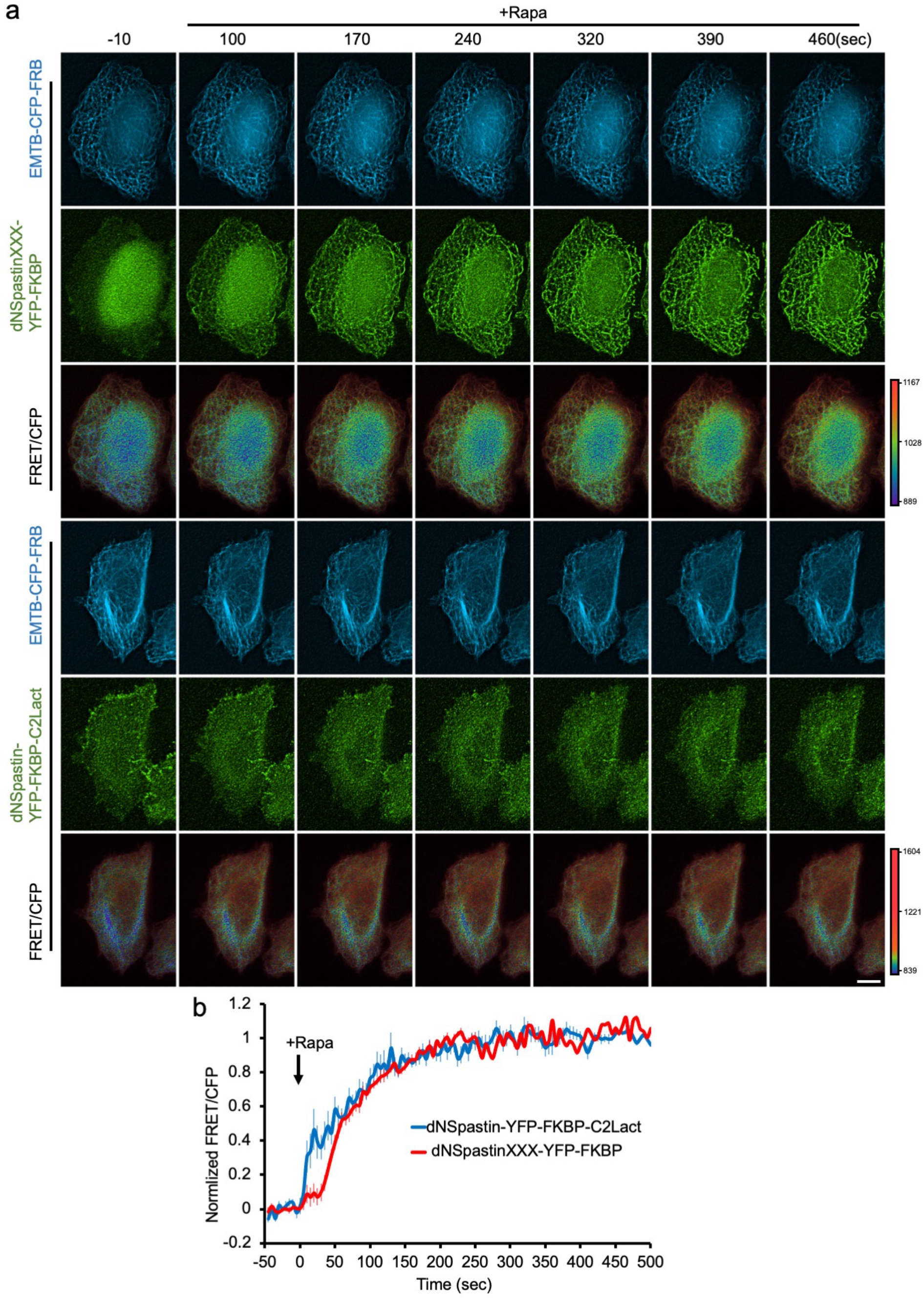
Rapidly recruiting engineered Spastins onto MTs. **a**. HeLa cells co-transfected with EMTB-CFP-FRB (blue) and either dNSpastinXXX-YFP-FKBP or dNSpastin-YFP-FKBP-C2Lact (green) were treated with 100 nM rapamycin (Rapa). The FRET signal of cells was monitored in real time by live-cell imaging. Scale bar, 10 μm. **b**. The normalized intensity of FRET/CFP in cells upon rapamycin treatment. n = 23 and 25 cells for dNSpastinXXX-YFP-FKBP (red) and dNSpastin-YFP-FKBP-C2Lact (blue), respectively, from four independent experiments. Data are shown as the mean ± S.E.M.

**Supplementary Fig. 4.**
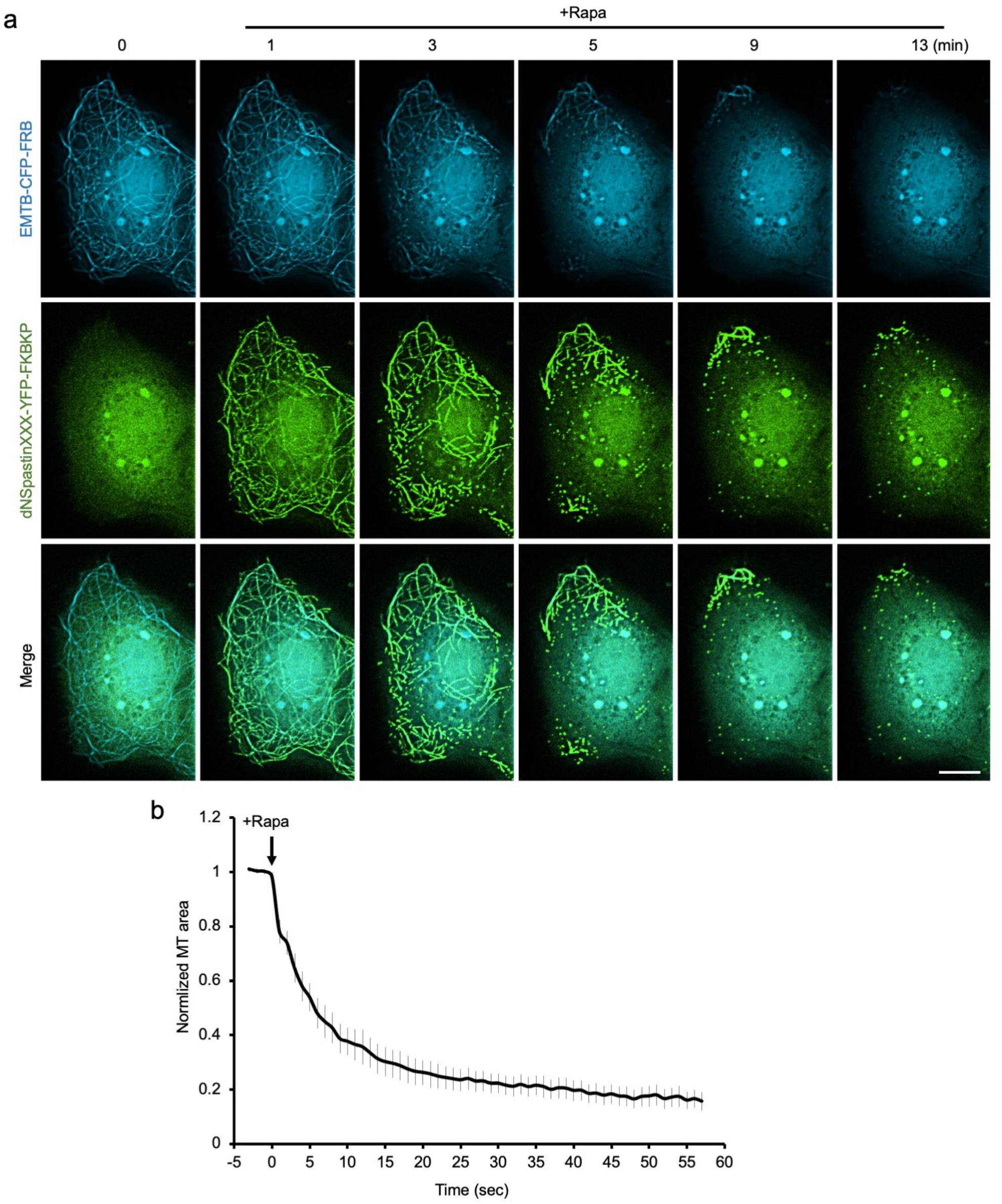
Rapid disruption of MTs in COS7 cells. **a**. COS7 cells transfected with dNSpastinXXX-YFP-FKBP (green) and EMTB-CFP-FRB (blue) were treated with rapamycin (100 nM) and imaged. Scale bar, 10 μm. **b**. The normalized MT filament area in cells transfected and treated as in **a**. n = 12 cells from four independent experiments. Data are shown as the mean ± S.E.M.

**Supplementary Fig. 5.**
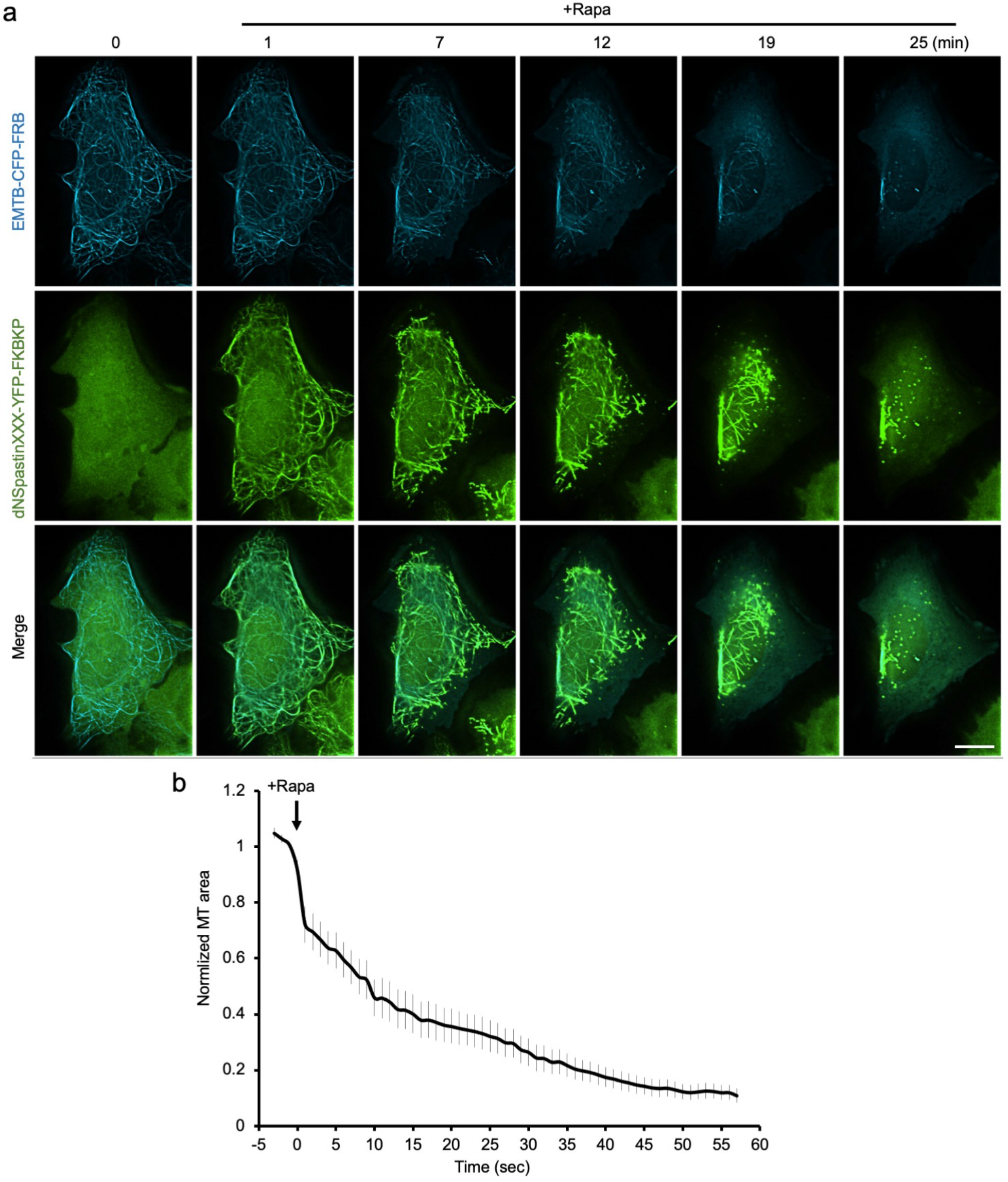
Rapid disruption of MTs in U2OS cells. **a**. U2OS cells transfected with dNSpastinXXX-YFP-FKBP (green) and EMTB-CFP-FRB (blue) were treated with rapamycin (100 nM) and imaged. Scale bar, 10 μm. **b**. The normalized MT filament area in cells transfected and treated as in **a**. n = 16 cells from four independent experiments. Data are shown as the mean ± S.E.M.

**Supplementary Fig. 6.**
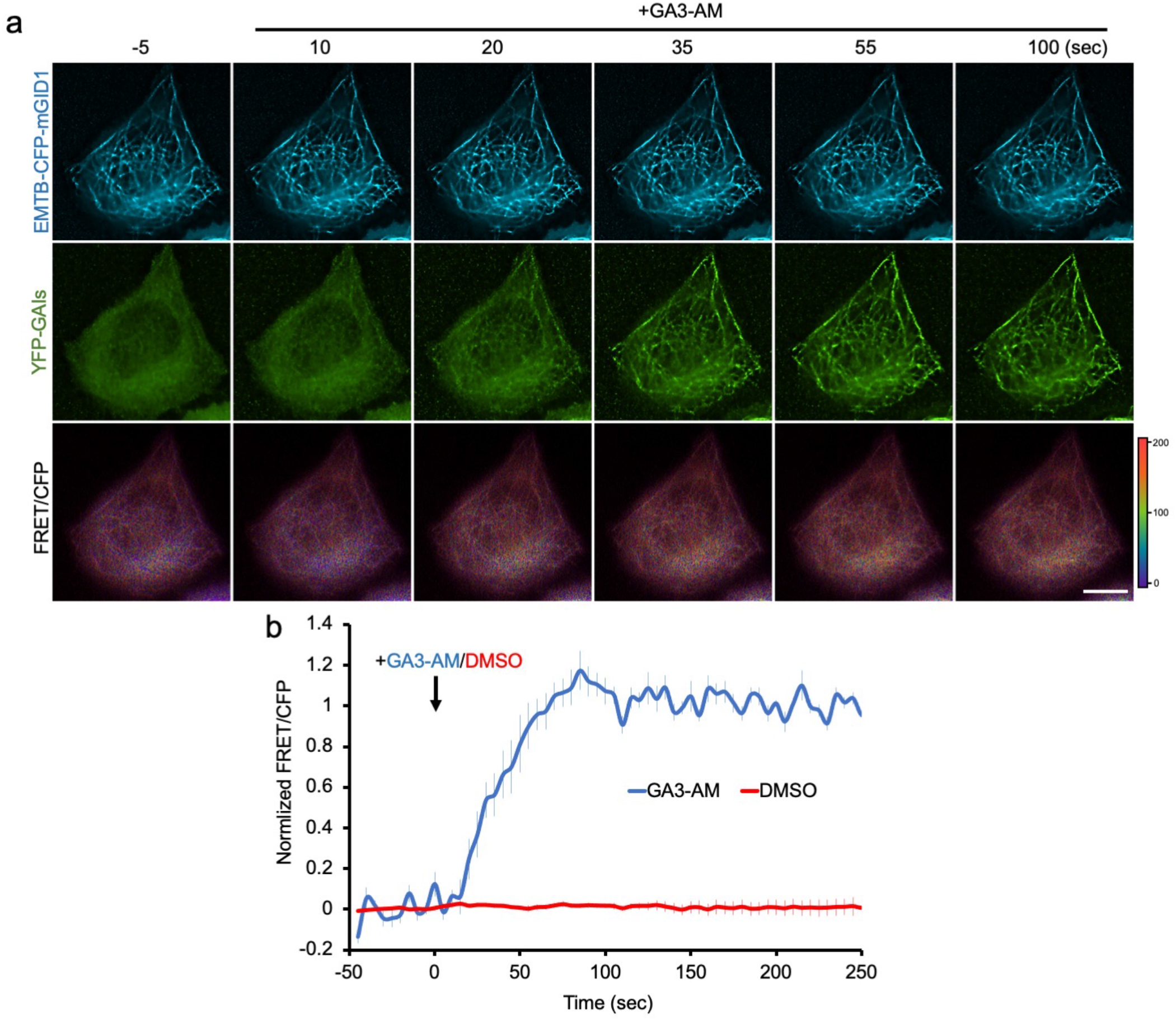
Using a gibberellin-based system to rapidly translocate proteins of interest onto MTs. **a**. HeLa cells co-transfected with EMTB-CFP-mGID1 (blue) and YFP-GAIs (green) (mGID1 and GAIs are dimerizing partners of the gibberellin system) were treated with GA3-AM (chemical dimerizer of the gibberellin system; 100 μM). We used live-cell imaging to monitor the FRET/CFP signal in cells. Scale bar, 10 μm. **b**. The normalized ratio of FRET/CFP in cells upon GA3-AM (blue) and 0.1% DMSO (red) treatment. n = 10 and 6 cells for the DMSO and GA3-AM treatments, respectively, from three independent experiments. Data are shown as the mean ± S.E.M.

**Supplementary Fig. 7.**
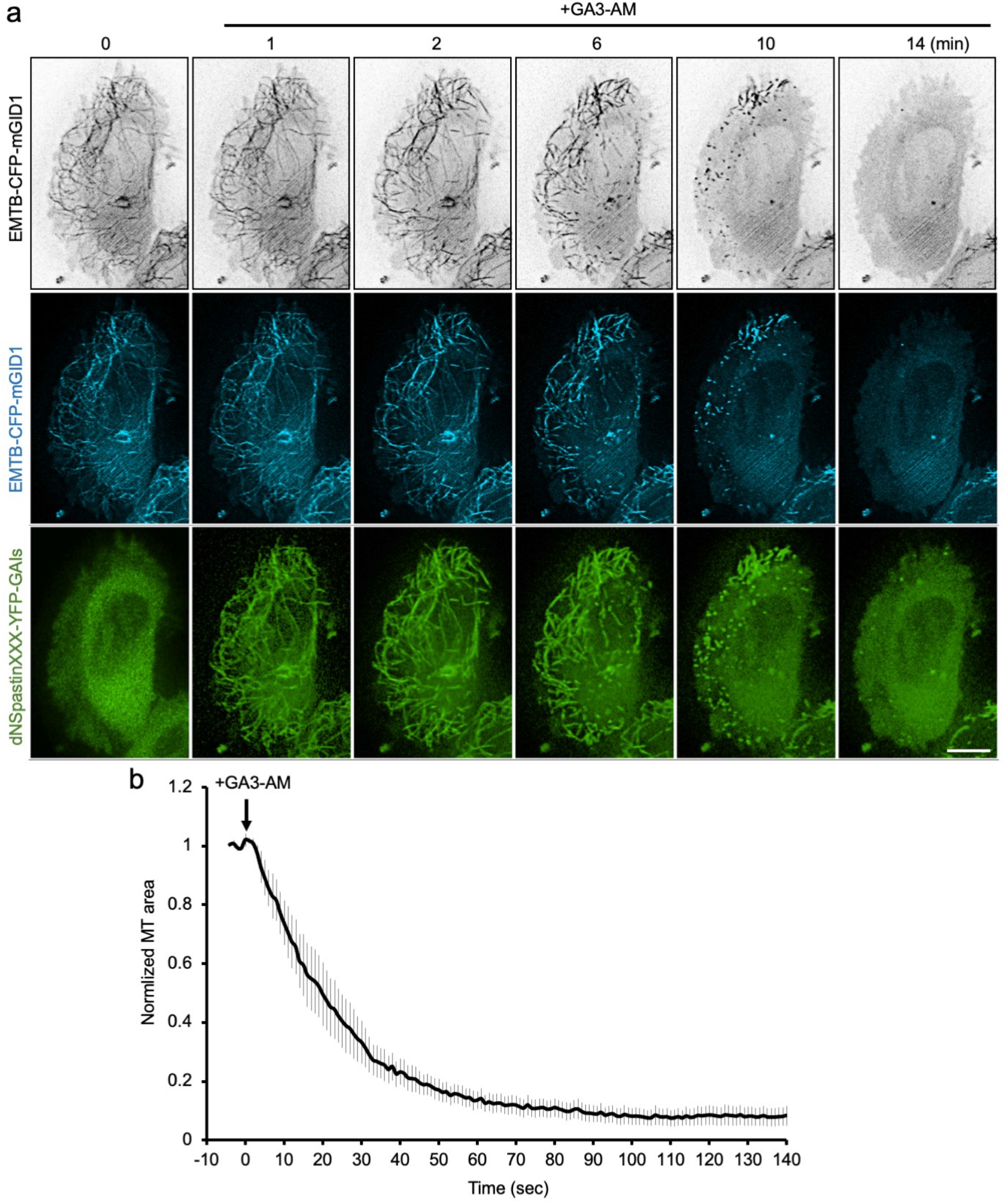
Using the gibberellin-based system to rapidly disassemble MTs. **a**. HeLa cells co-transfected with dNSpastinXXX-YFP-GAIs (green) and EMTB-CFP-mGID1 (black and blue) were treated with GA3-AM (100 μM). Scale bar, 10 μm. **b**. The normalized MT filament area in cells transfected with the constructs shown in **a** upon GA3-AM treatment. n = 8 cells from three independent experiments. Data are shown as the mean ± S.E.M.

**Supplementary Fig. 8.**
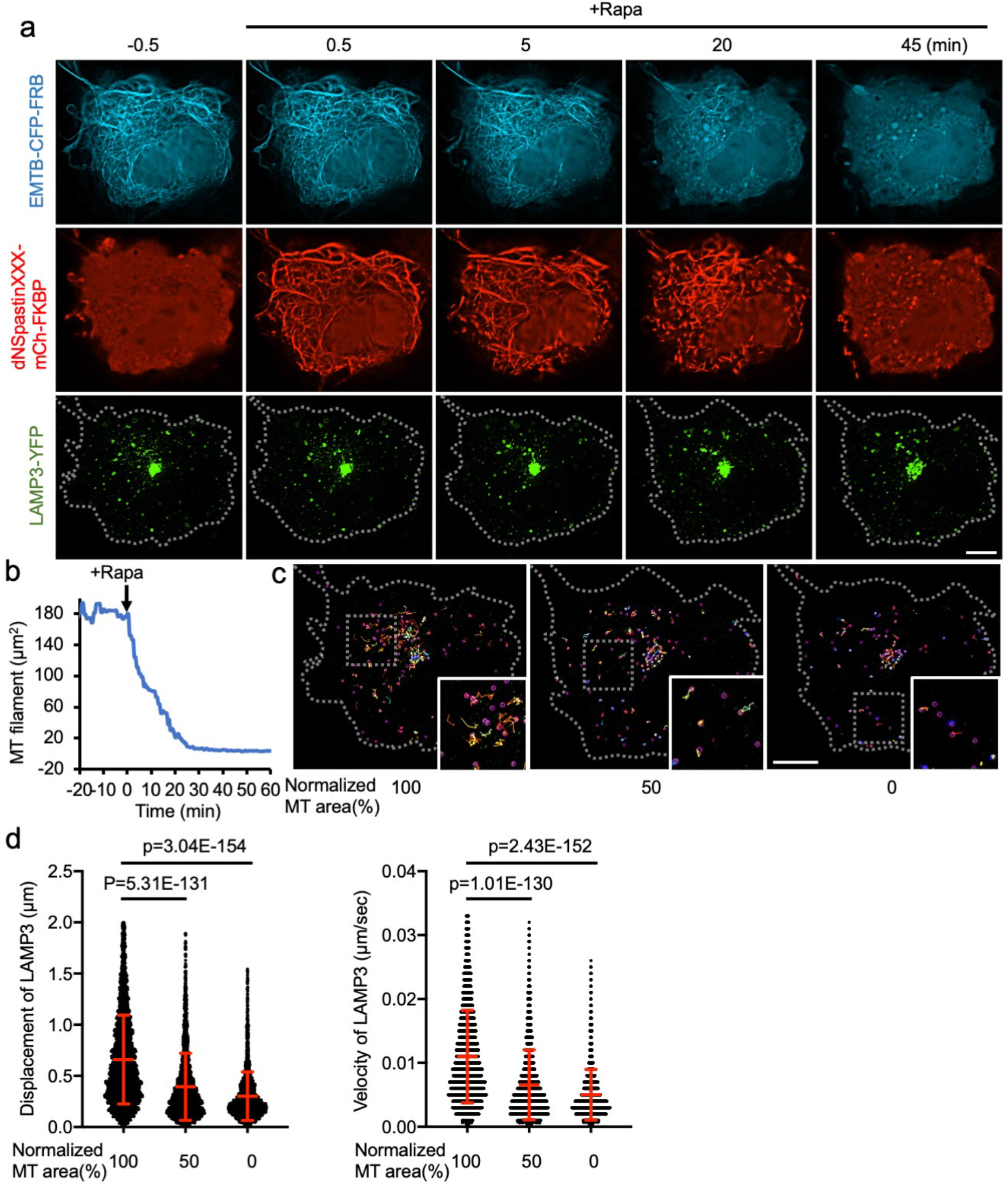
Acute MT disassembly attenuates lysosome dynamics. **a**. HeLa cells co-transfected with EMTB-CFP-FRB (blue), dNSpastinXXX-mCh-FKBP (red), and LAMP3-YFP (green) were treated with rapamycin (100 nM) to induce MT disruption. Dotted lines indicate the cell boundary. Scale bar, 10 μm. **b**. The MT filament area in the cell shown in **a. c**. Trajectories of each LAMP3-YFP−labeled lysosome at different levels of MT disruption. Insets show higher-magnification images of the regions indicated by the dotted-line boxes. Scale bar, 10 μm. **d**. Displacement (left) and velocity (right) of each labeled lysosome shown in **c**. n = 2893, 2618, and 2315 lysosomes in the MT 100, 50, and 0% group, respectively. Data (red) are shown as the mean ± SD. Student’s t-tests were performed to generate the indicated p-values.

**Supplementary Fig. 9.**
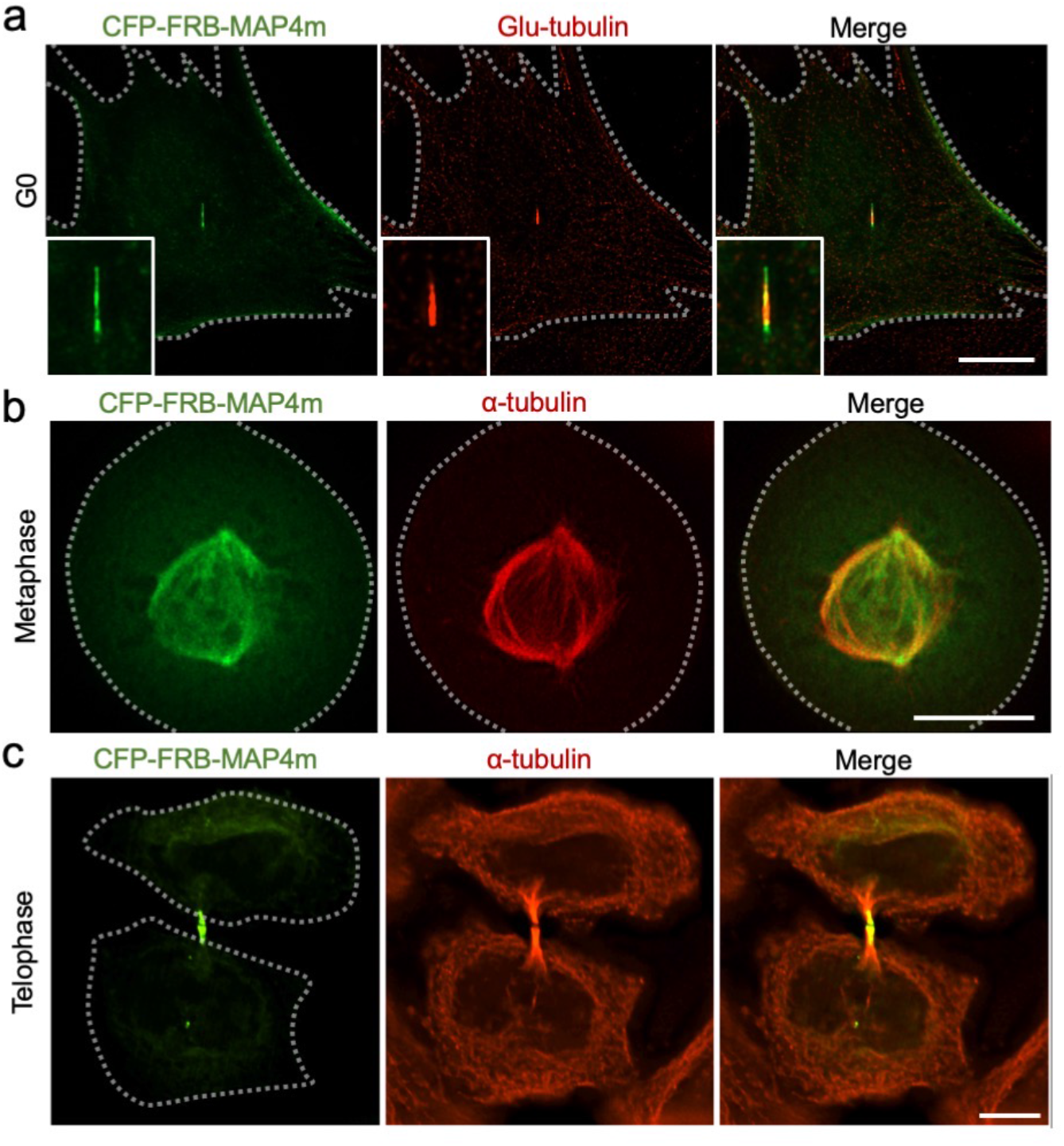
Subcellular distribution of CFP-FRB-MAP4m at different phases of the cell cycle. **a**. NIH3T3 cells transfected with CFP-FRB-MAP4m (green) were serum starved to promote ciliogenesis. Ciliated cells were fixed and labeled with GT335, an antibody that is specific for glutamylated tubulin (Glu-tub; red) at ciliary axonemes. **b** and **c**. HeLa cells expressing CFP-FRB-MAP4m (green) were synchronized during metaphase (**b**) and telophase (**c**) and then were fixed and stained with α-tubulin antibody (red). Dotted lines indicate cell boundaries. Scale bar, 10 μm.

**Supplementary Fig. 10.**
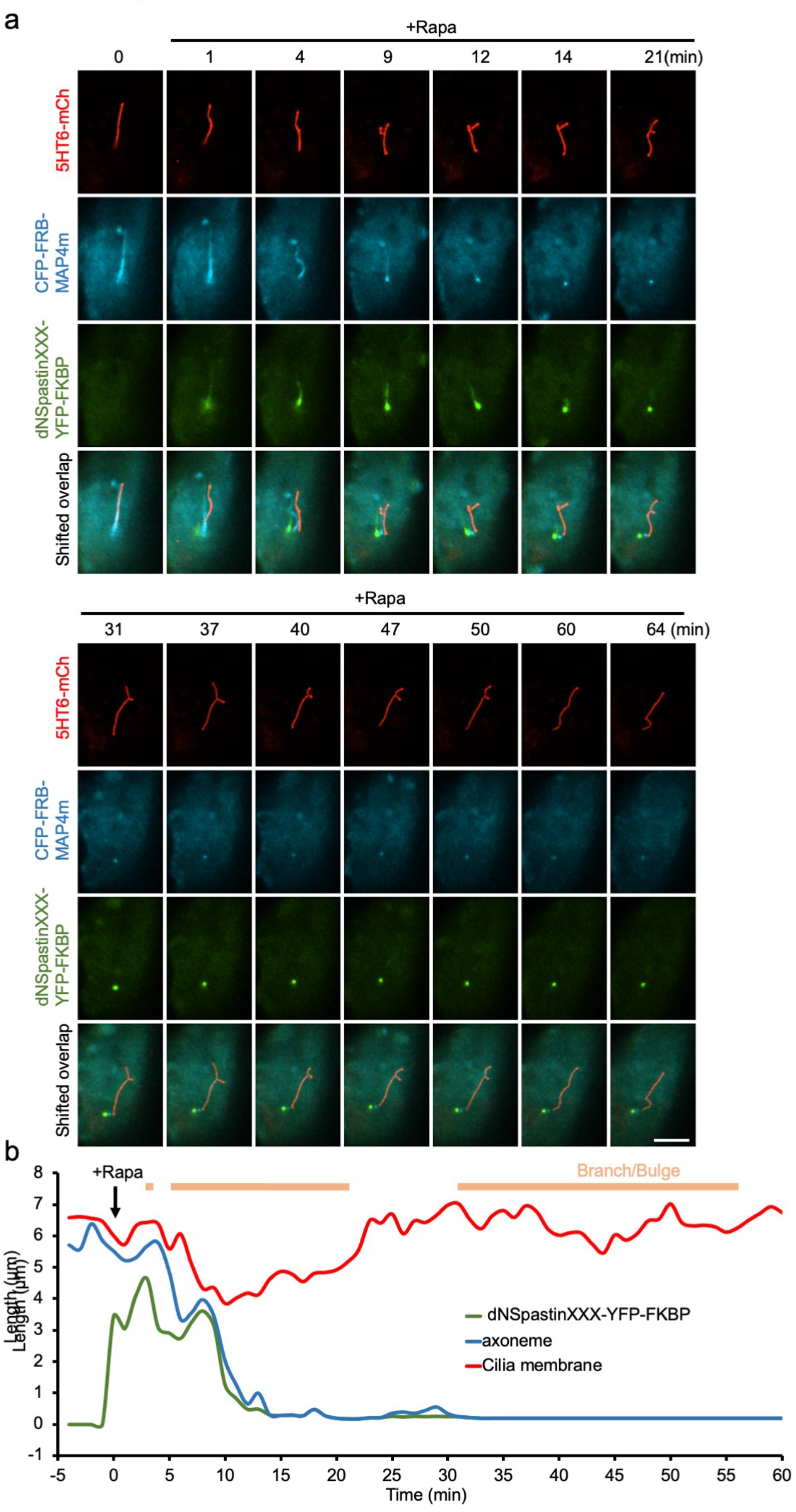
The structure of the cilia occasionally does not collapse after axoneme disruption. **a**. NIH3T3 cells co-transfected with 5HT6-mCh (ciliary membrane; red), CFP-FRB-MAP4m (axonemal MTs; blue), and dNSpastinXXX-YFP-FKBP (green) were serum starved to induce ciliogenesis. Ciliated cells were treated with rapamycin (100 nM) to recruit dNSpastinXXX-YFP-FKBP onto axonemes. The recruitment of Spastin proteins, morphology of the axonemes, and ciliary membrane were monitored upon rapamycin addition by live-cell imaging. Scale bar, 5 μm. **b**. The length of the indicated structures in the cell shown in **a** was measured and plotted. The time period during which the ciliary membrane became bulged and/or branched is shown with the orange lines above the graph.

**Supplementary Fig. 11.**
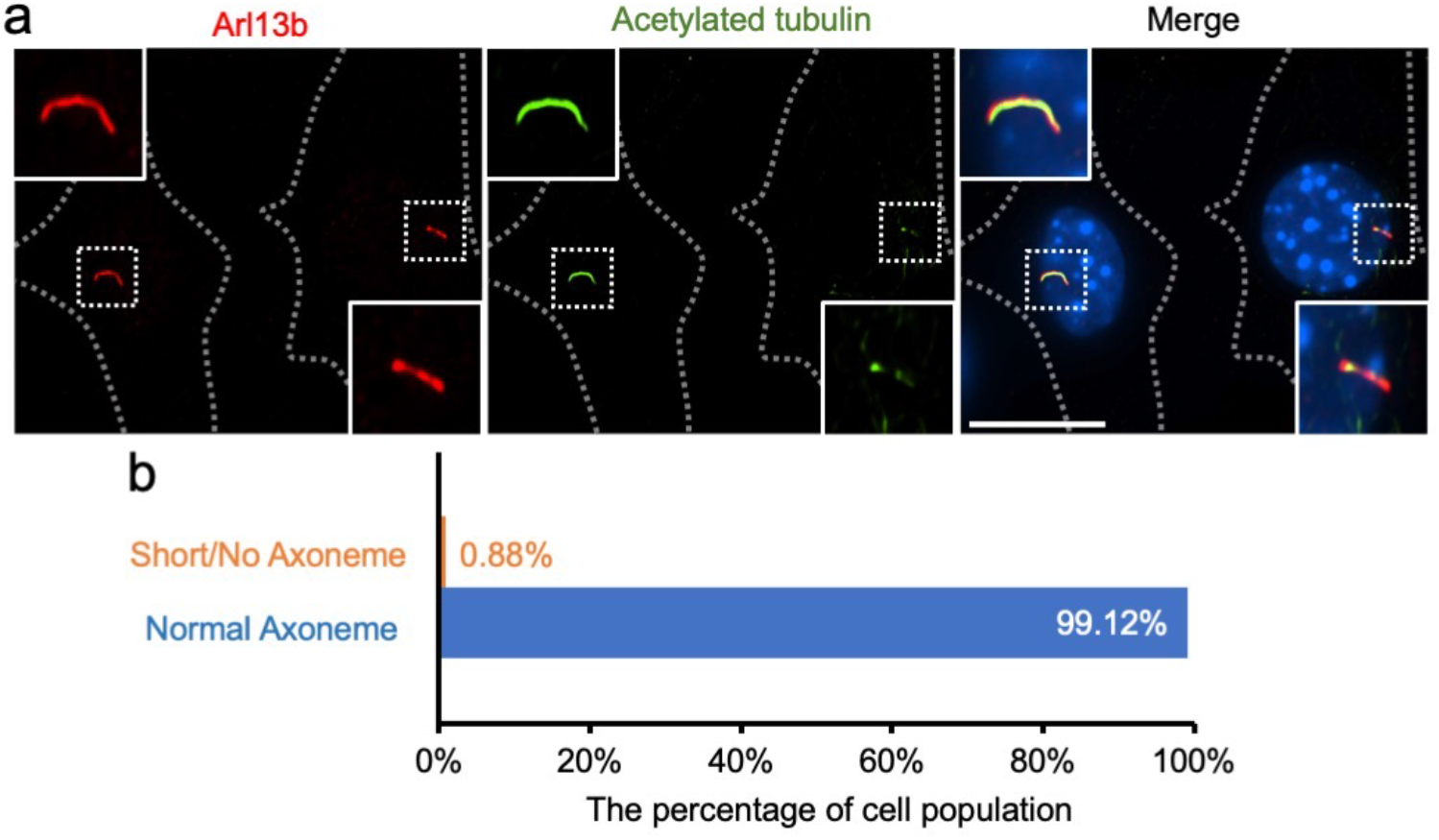
Cells can spontaneously form ciliary membranes without axonemes. **a**. NIH3T3 cells were serum starved to promote ciliogenesis. Ciliated cells were fixed and labeled with antibodies against Arl13b (a ciliary membrane marker; red) and against acetylated tubulin (acetylated tub; a ciliary axoneme marker; green). Dotted lines indicate cell boundaries. Insets show higher-magnification images of the regions indicated by the dotted-line boxes. Scale bar, 10 μm. **b**. The percentage of cells that spontaneously formed ciliary membranes with a short or absent axoneme or a normal axoneme. n = 391 cells from four dishes.

**Supplementary Fig. 12.**
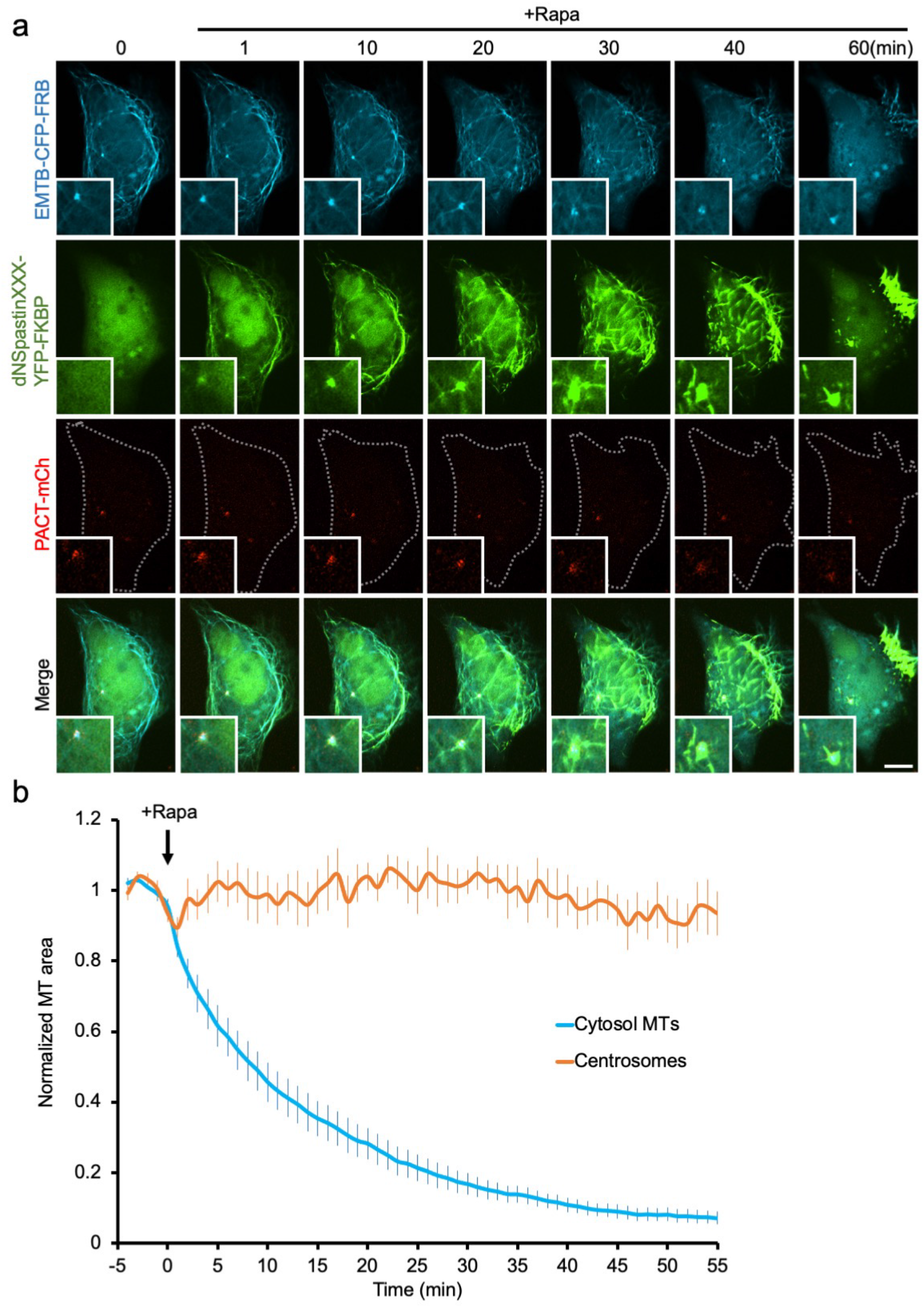
Spastin does not disrupt centrosomes. **a**. HeLa cells co-transfected with EMTB-CFP-FRB (blue), dNSpastinXXX-YFP-FKBP (green), and PACT-mCh (a centrosomal marker; red) were treated with rapamycin (100 nM) and imaged. Dotted lines indicate the cell boundary. Insets show higher-magnification images of the centrosome regions. Scale bar, 10 μm. **b**. The normalized area of MT filaments in the cytosol (blue) and centrosomes (orange) is shown. n = 8 cells from four independent experiments. Data are shown as the mean ± S.E.M.

**Supplementary Fig. 13.**
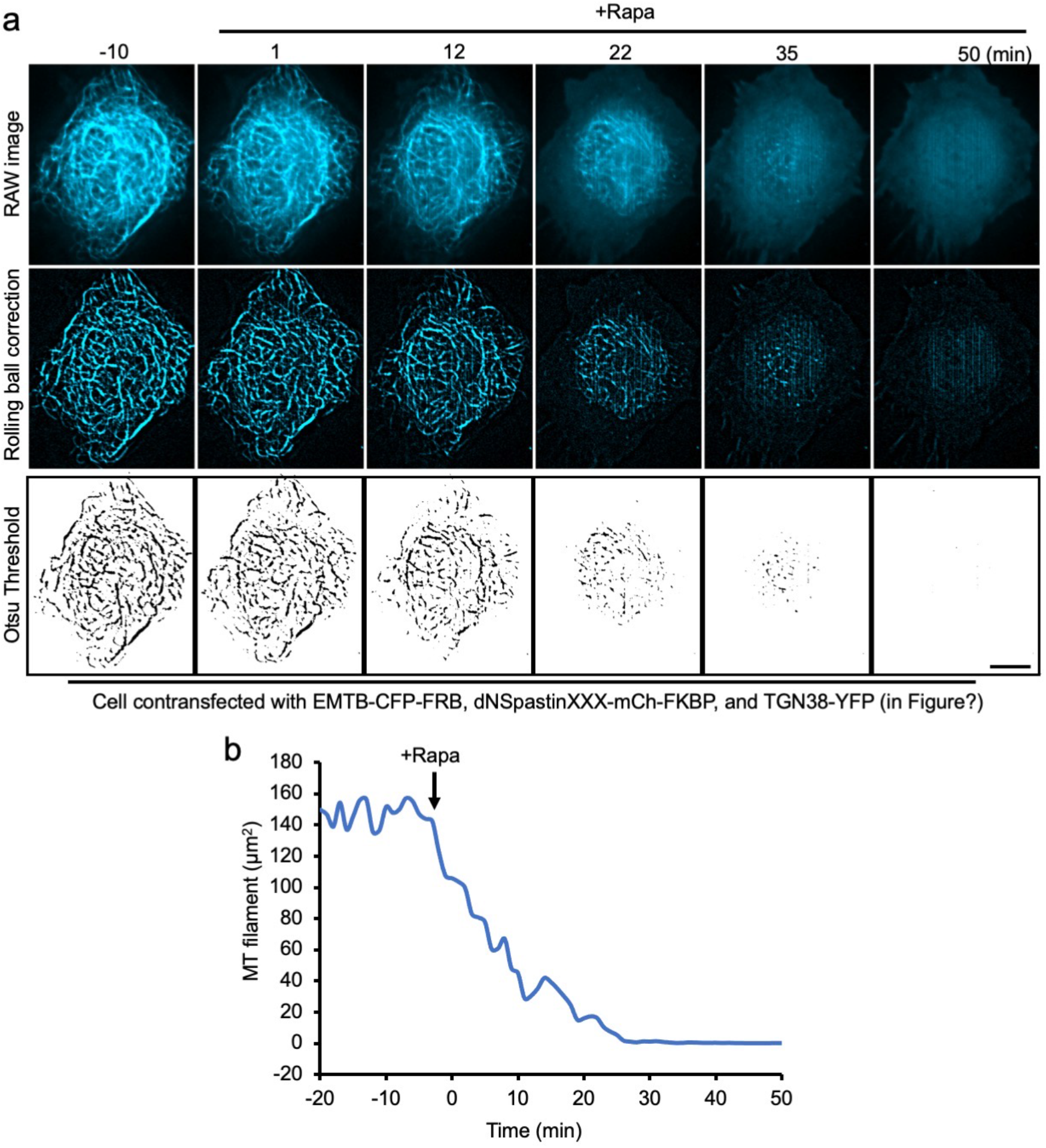
Method used to measure MT filament area in real time. **a**. MT filaments in living cells were labeled with EMTB-CFP-FRB and used for real-time imaging (RAW image). The cytosol background of the RAW images was removed by rolling ball correction and processed to generate the binary MT filament pattern via the Otsu threshold. Scale bar, 10 μm. **b**. The area of binary MT filaments upon MT disruption treatment was quantified and plotted.

**Extended data Fig. 1.**
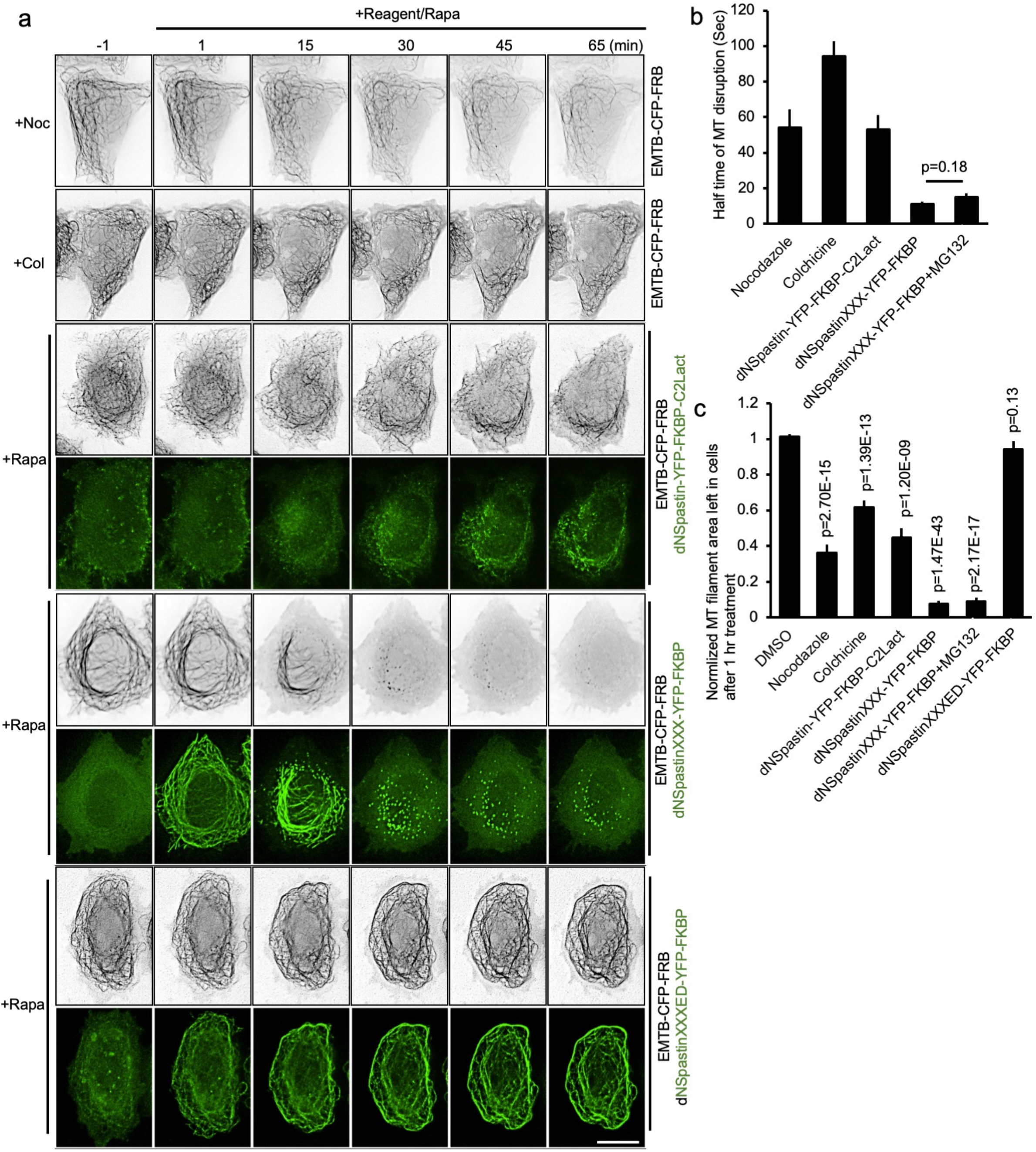
The MT disruption rates of MTAs and our MT disruption system. **a**. HeLa cells transfected with the indicated constructs were treated with nocodazole (10 μM), colchicine (500 μM), or rapamycin (100 nM). Scale bar, 10 μm. **b**. The half time of MT disassembly triggered by the indicated MT disruption systems. n = 30, 45, 19, 30, and 6 cells from left to right, three to five independent experiments. **c**. The relative proportion of MT area left in transfected HeLa cells after 1 hr of different MT-disrupting treatments. n = 34, 30, 45, 19, 20, 6, and 14 cells from left to right, three to five independent experiments. Data are shown as the mean ± S.E.M. Student’s t-tests were performed, with the resulting p-values indicated.

**Extended data Fig. 2.**
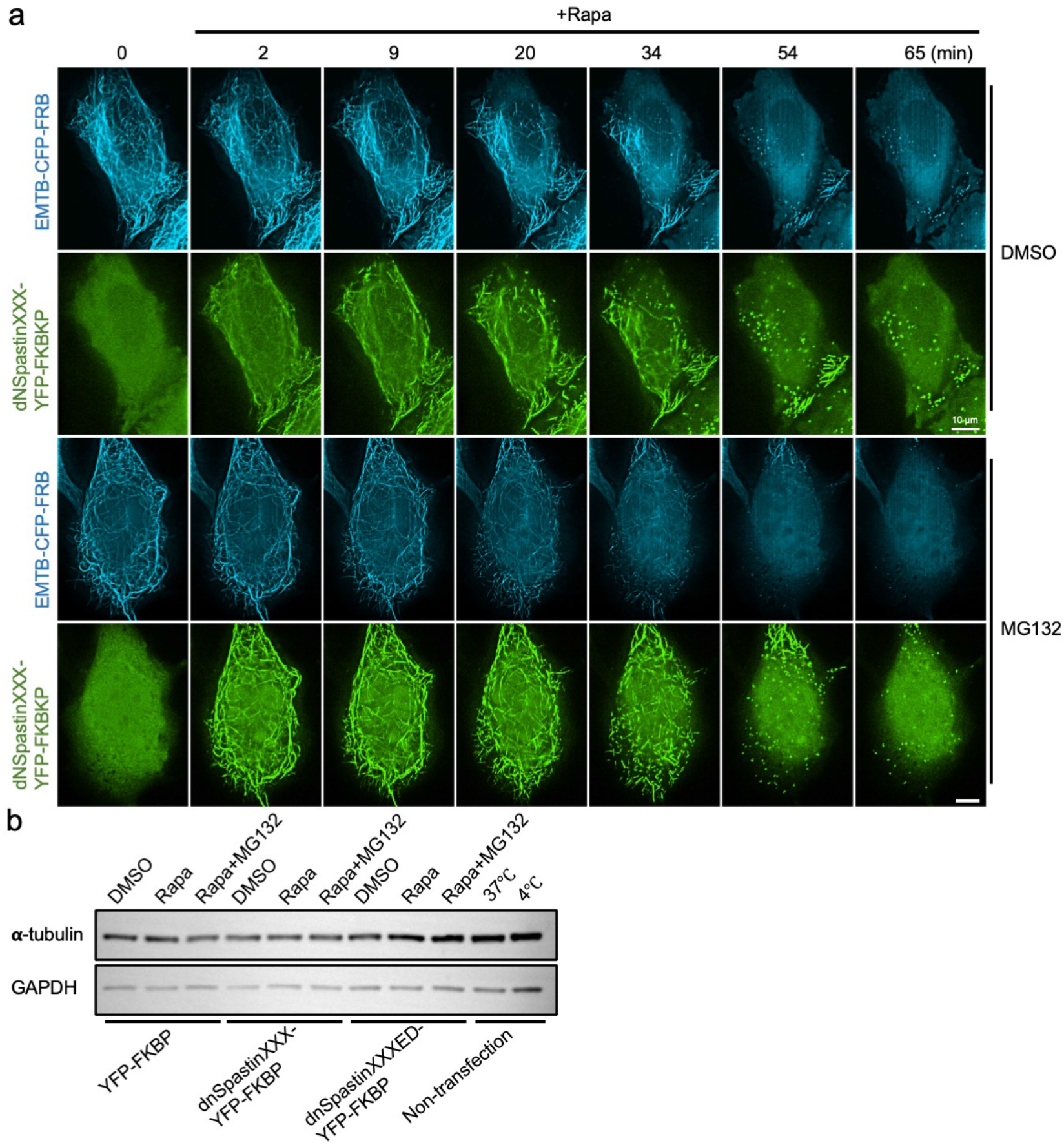
Acute MT disruption is not dependent on proteasome-mediated degradation. **a**. COS7 cells co-expressing EMTB-CFP-FRB (blue) and dNSpastinXXX-YFP-FKBP (green) were either pre-treated with 0.1% DMSO (control) or proteasome inhibitor, MG132 (50 μM) for 30 mins. The effect of MG132 on dNSpastinXXX-mediated MT disruption was evaluated by live-cell imaging. Scale bar, 10 μm. **b**. The α-tubulin protein levels in HeLa cells co-transfected with EMTB-CFP-FRB and the indicated constructs after 30 min of DMSO or MG132 (50 μM) pretreatment followed by rapamycin (100 nM) treatment for the indicated time. Cold-mediated MT disassembly at 4°C was included as a control.

**Extended data Fig. 3.**
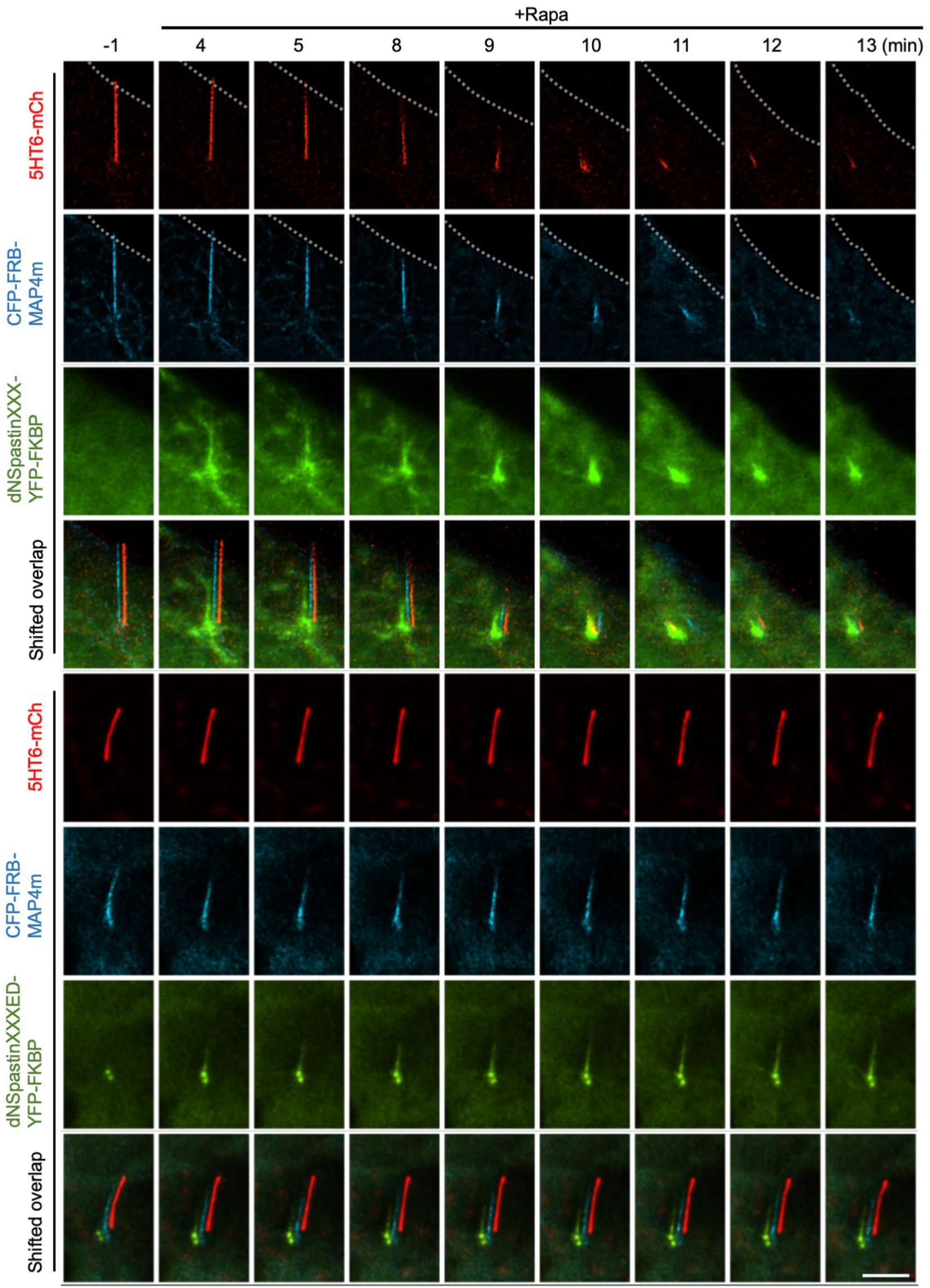
Rapid disassembly of primary cilia. **a**. NIH3T3 cells co-transfected with 5HT6-mCh (a ciliary membrane marker; red), CFP-FRB-MAP4m (an axoneme marker; blue), and dNSpastinXXX-YFP-FKBP (upper panel; green) or enzyme dead dNSpastinXXXED-YFP-FKBP (lower panel; green) were serum starved for 24 hr to induce ciliogenesis. Ciliated cells were treated with rapamycin (100 nM) to recruit dNSpastin proteins onto axonemes. The recruitment of Spastin proteins, morphology of ciliary axonemes, and ciliary membrane, upon rapamycin treatment was monitored by live-cell imaging. Dotted lines indicate the cell boundary. Scale bar, 5 μm.

**Extended data Fig. 4.**
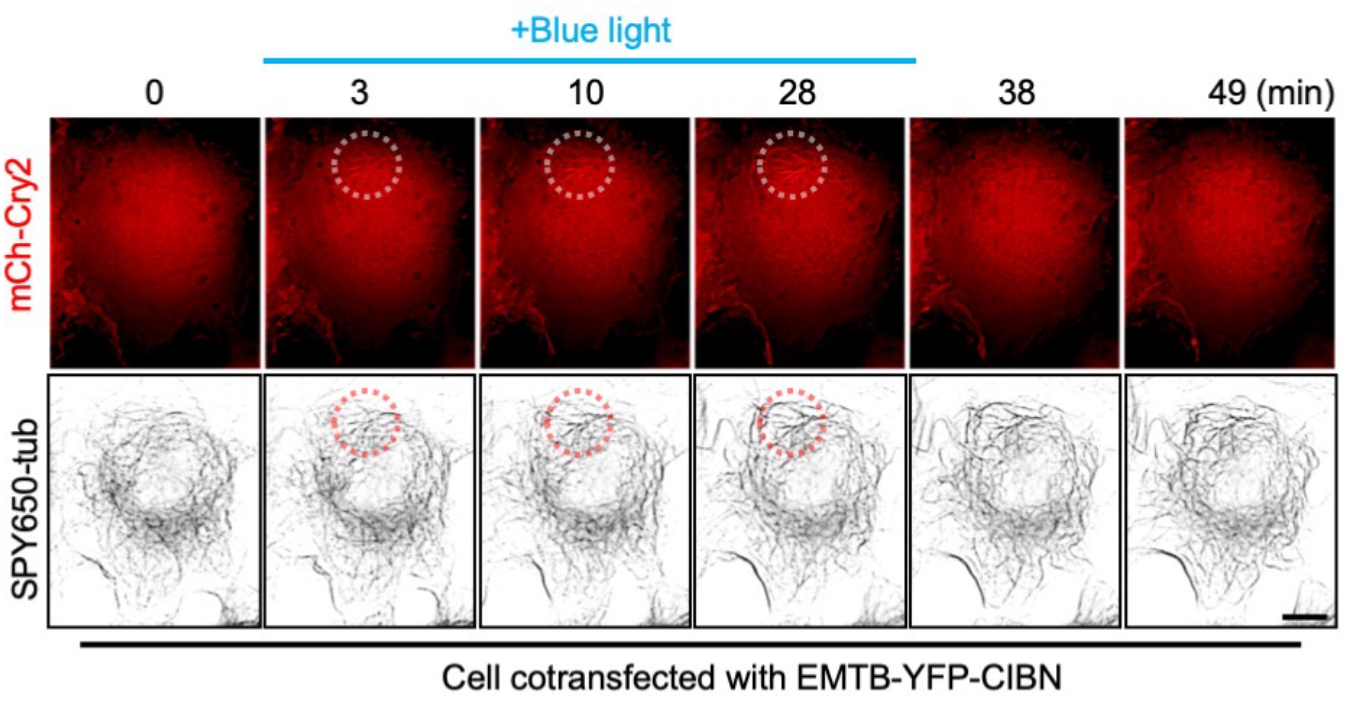
Recruitment of light-sensitive dimerizing proteins without spastin does not disrupt MTs. COS7 cells were co-transfected with EMTB-YFP-CIBN and mCh-Cry2 (red), and MTs were labeled with SPY650-tubulin (SPY650-tub; black). Illumination with blue light for the indicated period of time triggered rapid recruitment of mCh-Cry2 onto MTs in the light-illuminated region (dotted circle). The MTs, however, remained intact after recruitment of mCh-Cry2. Scale bar, 10 μm.

**Supplementary Video 1: Rapid translocation of POIs onto MTs**. HeLa cells were co-transfected with EMTB-CFP-FRB (blue) and YFP-FKBP (green) 1 day before imaging. The addition of 100 nM rapamycin led to a rapid increase in YFP-FKBP fluorescence and the FRET/CFP signal (heatmap) at MTs. Images were taken every 5 sec for 3 min. Scale bar, 10 μm. See also Fig. 1b.

**Supplementary Video 2: Rapid recruitment of engineered Spastins onto MTs**. HeLa cells were co-transfected with EMTB-CFP-FRB (blue) and either dNSpastinXXX-YFP-FKBP (upper panel; green) or dNSpastin-YFP-FKBP-C2Lact (lower panel; green). The addition of 100 nM rapamycin led to a rapid increase in YFP-FKBP fluorescence and FRET/CFP (heatmap) at MTs. Images were taken every 10 sec for 10 min. Scale bar, 10 μm. See also Supplementary Fig. 3a.

**Supplementary Video 3: Slow MT disruption in the presence of two MTAs**. HeLa cells transfected with EMTB-CFP-FRB were treated with either nocodazole (10 μM; left) or colchicine (500 μM; right). Images were taken every 1 min for 70 min. Scale bar, 10 μm. See also Extended data Fig. 1a.

**Supplementary Video 4: Rapid MT disruption with our MT disruption system**. HeLa cells co-transfected with EMTB-CFP-FRB (black) and the indicated Spastin constructs (green) were treated with rapamycin (100 nM). Images were taken every 1 min for 70 min. Scale bar, 10 μm. See also Fig. 1d and Extended data Fig. 1a.

**Supplementary Video 5: Rapid disruption of MTs in COS7 cells**. COS7 cells transfected with dNSpastinXXX-YFP-FKBP (green) and EMTB-CFP-FRB (blue) were treated with rapamycin (100 nM) and imaged. Images were taken every 1 min for 1 hr. Scale bar, 10 μm. See also Supplementary Fig. 4a.

**Supplementary Video 6: Rapid disruption of MTs in U2OS cells**. U2OS cells transfected with dNSpastinXXX-YFP-FKBP (green) and EMTB-CFP-FRB (blue) were treated with rapamycin (100 nM) and imaged. Images were taken every 1 min for 1 hr. Scale bar, 10 μm. See also Supplementary Fig. 5a.

**Supplementary Video 7: The gibberellin system rapidly disrupts MTs**. HeLa cells co-transfected with dNSpastinXXX-YFP-GAIs (green) and EMTB-CFP-mGID1 (blue and black) were treated with GA3-AM (100 μM). Images were taken every 1 min for 1 hr. Scale bar, 10 μm. See also Supplementary Fig. 7a.

**Supplementary Video 8: Acute MT disruption is independent of proteasome-mediated degradation**. COS7 cells co-expressing EMTB-CFP-FRB (blue) and dNSpastinXXX-YFP-FKBP (green) were either pretreated with 0.1% DMSO (upper panel) or the proteasome inhibitor MG132 (50 μM) for 30 min (lower panel). Rapamycin (100 nM)-mediated recruitment of dNSpastinXXX led to a similar degree of MT disassembly after each pretreatment. Images were taken every 1 min for 90 min. Scale bar, 10 μm. See also Extended data Fig. 2a.

**Supplementary Video 9: Inhibition of Spastin activity reverses the MT disruption**. HeLa cells co-transfected with EMTB-CFP-FRB (blue and black) and dNSpastinXXX-YFP-FKBP (green) were treated with rapamycin (100 nM) for 28 min to induce acute MT disassembly followed by spastazoline (10 μM) treatment to halt our MT disruption system. Images were taken every 1 min for 2 hr. Scale bar, 10 μm. See also Fig. 2a.

**Supplementary Video 10: Acute MT disassembly impairs vesicular trafficking**. HeLa cells co-transfected with EMTB-CFP-FRB (blue), dNSpastinXXX-mCh-FKBP (red), and TGN38-YFP (green) were treated with rapamycin (100 nM). Images were taken every 1 min for 2 hr. Scale bar, 10 μm. See also Fig. 3.

**Supplementary Video 11: Trajectories of each labeled vesicle at different MT levels**. HeLa cells co-transfected with EMTB-CFP-FRB (blue), dNSpastinXXX-mCh-FKBP (red), and TGN38-YFP (green) were treated with rapamycin (100 nM). Trajectories of each labeled vesicle at normalized MT areas of 100% (upper panel), 50% (middle), and 0% (lower panel) are shown. Scale bar, 10 μm. See also Fig. 3c.

**Supplementary Video 12: Acute MT disassembly impairs lysosome dynamics**. HeLa cells co-transfected with EMTB-CFP-FRB (blue), dNSpastinXXX-mCh-FKBP (red), and LAMP3-YFP (green) were treated with rapamycin (100 nM). Images were taken every 30 sec for 90 min. Scale bar, 10 μm. See also Supplementary Fig. 8.

**Supplementary Video 13: Trajectories of each labeled lysosome at different levels of MT disruption**. HeLa cells co-transfected with EMTB-CFP-FRB (blue), dNSpastinXXX-mCh-FKBP (red), and LAMP3-YFP (green) were treated with rapamycin (100 nM). Trajectories of **each labeled lysosome** at normalized MT areas of 100% (upper panel), 50% (middle), and 0% (lower panel) are shown. Scale bar, 10 μm. See also Supplementary Fig. 8.

**Supplementary Video 14: Rapid translocation of POIs onto tyrosinated MTs**. COS7 cells were co-transfected with TagRFP-FRB-A1AY1 (red) and TagCFP-FKBP (green) 1 day before imaging. The addition of 100 nM rapamycin led to a rapid increase in TagCFP-FKBP fluorescence at A1AY1-labeled MTs. Images were taken every 30 sec for 14 min. Scale bar, 10 μm. See also Fig. 4c.

**Supplementary Video 15: Rapid disassembly of primary cilia**. NIH3T3 cells co-transfected with 5HT6-mCh (a marker of ciliary membranes; red), CFP-FRB-MAP4m (an axoneme marker; blue), and dNSpastinXXX-YFP-FKBP (upper panel; green) or enzyme dead dNSpastinXXXED-YFP-FKBP (lower panel; green) were serum starved for 24 hr to induce ciliogenesis, followed by treatment with rapamycin (100 nM) to recruit dNSpastin proteins onto axonemes. Images were taken every 1 min for 90 min. Scale bar, 5 μm. See also Fig. 5a and Extended data Fig. 3.

**Supplementary Video 16: Ciliary structures occasionally do not collapse after axoneme disruption**. NIH3T3 cells co-transfected with 5HT6-mCh (a marker of ciliary membranes; red), CFP-FRB-MAP4m (an axoneme marker; blue), and dNSpastinXXX-YFP-FKBP (green) were serum starved to induce ciliogenesis. Ciliated cells were treated with rapamycin (100 nM) to recruit dNSpastin proteins onto axonemes. Images were taken every 1 min for 90 min. Scale bar, 5 μm. See also Supplementary Fig. 10a.

**Supplementary Video 17: Mitotic spindles can be rapidly disrupted in metaphase cells**. HeLa cells co-transfected with H2B-mCherry (a chromosomal marker; red), CFP-FRB-MAP4m (a marker of mitotic spindles; blue), and dNSpastinXXX-YFP-FKBP (green) were synchronized in metaphase and treated with rapamycin (100 nM), which led to dNSpastinXXX-YFP-FKBP recruitment at mitotic spindles. Images were taken every 1 min for 90 min. Scale bar, 10 μm. See also Fig. 5c.

**Supplementary Video 18: Intercellular bridges can be rapidly disrupted in telophase cells**. HeLa cells co-transfected with H2B-mCherry (a chromosomal marker; red), CFP-FRB-MAP4m (a marker of intercellular bridges; blue), and dNSpastinXXX-YFP-FKBP (green) were treated with rapamycin (100 nM), which led to dNSpastinXXX-YFP-FKBP recruitment at intercellular bridges. Images were taken every 1 min for 1 hr. Scale bar, 10 μm. See also Fig. 5e.

**Supplementary Video 19: Centrosomes remain intact after acute MT disassembly triggered by dNSpastinXXX**. HeLa cells co-transfected with EMTB-CFP-FRB (blue), dNSpastinXXX-YFP-FKBP (green), and PACT-mCh (a centrosomal marker; red) were treated with rapamycin (100 nM). Images were taken every 1 min for 90 min. Scale bar, 10 μm. See also Supplementary Fig. 12a.

**Supplementary Video 20: A light-induced system leads to rapid translocation of the proteins of interest onto MTs**. COS7 cells co-transfected with EMTB-YFP-CIBN and mCh-Cry2 (red) were transiently illuminated by blue light in the regions indicated by the dotted circles. MTs were labeled by SPY650-tubulin (black). Illumination rapidly triggered translocation of mCh-Cry2 onto MTs only within the illuminated region. Images were taken every 15 sec for 15 min. Scale bar, 10 μm. See also Fig. 6a.

**Supplementary Video 21: A light-induced system] leads to spatially and temporally specific disassembly of MTs**. COS7 cells co-transfected with EMTB-YFP-CIBN and dNSpastinXXX-mCh-Cry2 (red) were transiently illuminated by blue light in the regions indicated by the dotted circles. MTs were labeled by SPY650-tubulin (green). The light illumination rapidly triggered MT disruption in the illuminated regions (dotted circle). MT regrowth occurred when the light was off. Images were taken every 1 min for 1 hr. Scale bar, 10 μm. See also Fig. 6c.

**Supplementary Video 22: Recruitment of light-sensitive dimerizing proteins without spastin does not disrupt MTs**. COS7 cells were co-transfected with EMTB-YFP-CIBN and mCh-Cry2 (red), and MTs were labeled by SPY650-tubulin (black). The cells were transiently illuminated by blue light in the regions indicated by the dotted circles. This triggered rapid recruitment of mCh-Cry2 onto MTs in illuminated regions. MTs remained intact after recruitment of mCh-Cry2. Images were taken every 1 min for 1 hr. Scale bar, 10 μm. See also Extended data Fig. 4.

## Methods

### Cell culture and transfection

COS7, HeLa, U2OS, HEK293T, and NIH3T3 cells were maintained at 37°C, 5% CO_2_, and 95% humidity in Dulbecco’s modified Eagle’s medium (DMEM) supplemented with 10% fetal bovine serum (FBS), and penicillin and streptomycin (Corning). To induce ciliogenesis, NIH3T3 cells were serum starved for 24 hr. COS7 cells were transfected with plasmid DNAs by using TurboFect transfection reagents (Thermo Fisher). HeLa, U2OS, HEK293T, and NIH3T3 cells were transfected with FuGENE HD (Promega). Transfected cells were incubated for 24−48 hr prior to imaging and other experiments.

### DNA constructs

We obtained constructs encoding the most abundantly expressed isoform of Spastin in cells (58 kDa; starting at position M85 in the mouse spastin sequence), a truncated form of dNSpastin that is missing the N-terminal 1−140 amino acids (the shortest active truncated version of spastin), and the enzyme-inactive versions SpastinFLED and dNSpastinED, each of which was tagged with EYFP (SpastinFL-YFP, dNSpastin-YFP, SpastinFLED-YFP, and dNSpastinED-YFP), from Dr. Carsten Janke (Institut Curie). To remove the MT binding domain of dNSpastin, the catalytic AAA domain of dNSpastin (dNSpstinCD) was amplified by PCR-based methods. Three residues of Spastin were mutated to XXX (dNSpastinXXX) by site-directed mutagenesis DNA encoding individual forms of dNSpastin (dNSpastin, dNSpastinED, dNSpastinCD, and dNSpastinXXX) was then cloned into the YFP-FKBP vector (pEGFP-C1 backbone) to generate dNSpastin-YFP-FKBP, dNSpastinCD-YFP-FKBP, and dNSpastinXXX-YFP-FKBP. Using this method, we also generated the enzyme-inactive dNSpastinXXXED-YFP-FKBP construct. dNSpastin-YFP-FKBP-C2Lact was generated by inserting the C2Lact sequence between the HindIII and BamHI restriction sites in dNSpastin-YFP-FKBP. The DNA fragments of A1AY1, TagRFP, and TagCFP were synthesized with codon optimization and subcloned to FRB and FKBP vectors. We transformed each construct into competent cells and isolated single clones for DNA purification. All DNA constructs were verified by DNA sequencing. The detailed protein sequences of constructs used in this study are provided in supplementary sequence.

### Immunofluorescence staining

Cells cultured in borosilicate glass Lab-Tek eight-well chambers (Nunc) were fixed in 4% paraformaldehyde (Electron Microscopy Sciences) at room temperature for 15 min. Fixed cells were permeabilized with 0.1% Triton X-100 and then incubated in blocking solution (phosphate-buffered saline with 2% bovine serum albumin) for 30 min at room temperature. To label cytosolic MTs, primary ciliary membrane, and axonemal MTs, cells were incubated for 1 hr at room temperature with mouse antibody against α-tubulin (1:500; Sigma Aldrich, T6199), rabbit antibody against Arl13b (1:500; Proteintech, 17711-1-AP), mouse antibody against glutamylated tubulin (1:100; Adipogen, AG-20B-0020-C100), mouse antibody against acetylated tubulin (1:500; Sigma Aldrich, T7451), rat antibody against tyrosinated tubulin (1:100; Sigma Aldrich, MAB1864), and mouse antibody against detyrosinated tubulin (1:100; MERCK, AB3201) each of which was diluted in blocking solution. Cells were then washed with PBS and incubated for 1 hr with appropriate secondary antibodies (1:1000 dilution; Thermo Fisher) at room temperature.

### Western blotting

HEK293T cells were co-transfected with EMTB-CFP-FRB and YFP-FKBP, dNSpastinXXX-YFP-FKBP, or dNSpastinXXXED-YFP-FKBP. Two days after transfection, cells were incubated with 50 μM MG132 (Sigma Aldrich) for 30 min and then treated with 100 nM rapamycin or 0.1% DMSO (vehicle control) for 30 min prior to cell collection. For the cold treatment (4°C), untransfected HEK293T cells were put on ice for 40 min to depolymerize MTs. Cells were lysed in RIPA lysis buffer (50 mM Tris-HCl, pH 7.6; 2 mM EGTA; 9% NaCl; 1% Triton X-100) containing protease inhibitors (Roche). Protein concentrations were measured with the Bio-Rad Protein Assay. Cell lysates were diluted with 2× Laemmli sample buffer (Bio-Rad) and boiled at 95°C for 10 min and then underwent western blotting. After the transfer process, the PVDF membranes (Bio-Rad) were incubated with blocking buffer (5% skim milk in Tris-buffered saline with Tween 20; TBST) for 1 hr at room temperature and then stained with primary antibodies against α-tubulin (1:1000; Sigma Aldrich, T6199) and GAPDH (1:5000; Cell Signaling, 2118), which were diluted with blocking buffer, overnight at 4°C. Membranes were washed with TBST and then were incubated with horseradish peroxidase−conjugated secondary antibodies diluted in blocking buffer (anti-rabbit, 1:10000; anti-mouse, 1:5000) for 1 hr at room temperature. The bioluminescence signal was detected with Amersham™ ECL Select™ (GE Healthcare), and blot images were acquired with an iBright™ FL1500 Instrument (Thermo Fisher).

### Live-cell imaging

Live transfected cells cultured on poly(D-lysine)-coated glass coverslips (Hecht Assistent) were treated with either 100 nM rapamycin for rapid induction of protein dimerization and translocation or MT-targeting agents (10 μM nocodazole or 500 μM colchicine) during imaging. Live-cell imaging was conducted on a Nikon T1 inverted fluorescence microscope (Nikon) with a 60× oil objective (Nikon), a Prime camera (Photometrics), and 37°C, 5% CO_2_ heat stage (Live Cell Instrument). Rapid recruitment of POIs was imaged at 5- or 10-sec intervals, whereas the process of MT disruption was imaged at 1-min intervals. Images were obtained using Nikon NIS-Elements AR software and processed with Huygens Deconvolution Software (Scientific Volume Imaging). Image analysis was mainly performed with Nikon NIS-Elements AR software.

### Photostimulation

COS7 cells were plated on poly(D-lysine)-coated coverslips and cultured in six-well plates (Thermo Scientific) for 48 hr. Before imaging, cells were incubated with SPY650-tubulin (1000-fold dilution; Spirochrome) at 37°C for 1 hr. Local photostimulation was carried out with a fluorescence microscope (Nikon) equipped with a digital micromirror device, polygon 400 (MIGHTEX), and a 488-nm light source. Cells were illuminated with the blue light (5 sec on/1 sec off; 1.6 nW/μm^2^) for the indicated duration. The mCherry and SPY650-tubulin were simultaneously imaged during photostimulation using Nikon element AR software.

### Measurement of MT filament area

The MT filaments in living cells were labeled with EMTB-CFP-FRB and imaged in real time (RAW image). The rolling ball correction was used to remove the cytosol background from the RAW images, which was carried out with the Nikon NIS-Elements AR software. Images were processed to generate the binary MT filament pattern via the Otsu threshold and analyzed by Fiji software (Supplementary Fig. 13).

### Tracking vesicles and lysosomes

The displacement and velocity of LAMP3-YFP/TGN38-YFP were tracked and analyzed by the DoG detector and Simple LAP tracker in the Fiji software plugin Trackmate. Estimated blob diameter was set at 0.8−1 μm, linking max distance at 2 microns, gap-closing distance at 2 μm, and gap-closing max frame gap at 0.

### Cell synchronization

Plasmid DNA transfection was carried out 20−24 hr prior to cell cycle synchronization. For synchronization, HeLa cells were treated with 2 mM thymidine (Sigma) for 16−18 hr to induce arrest at G1/S phase and then were treated with 2.5 ng/ml RO3306 (Sigma) for 12 hr to induce arrest at G2/M phase. After being washed with warm DMEM, cells were incubated with DMEM and 10% FBS at 37°C, 5% CO_2_ for 30−60 min to enrich the population of metaphase and telophase cells.

### Statistical analysis

We first determined whether variances were equal or not with the F-test and then used the unpaired two-tailed Student’s t-test to calculate p-values via PRISM 6 software. A p-value of <0.05 indicated a significant difference, and p < 0.01 indicated a highly significant difference.

